# NHA2 promotes cyst development in an *in vitro* model of polycystic kidney disease

**DOI:** 10.1101/364679

**Authors:** Hari Prasad, Donna K. Dang, Kalyan C. Kondapalli, Niranjana Natarajan, Valeriu Cebotaru, Rajini Rao

**Affiliations:** Department of Physiology, The Johns Hopkins University School of Medicine, Baltimore, MD; Department of Medicine, University of Maryland School of Medicine, Baltimore, MD

**Keywords:** Na^+^/H^+^ exchanger, polycystin, vasopressin, Ca^2+^ signaling, NFAT

## Abstract

Autosomal dominant polycystic kidney disease (ADPKD) is caused by mutations in *PKD1* and *PKD2* encoding polycystin-1 (PC1) and polycystin-2 (PC2), respectively. The molecular pathways linking polycystins to cyst development in ADPKD are still unclear. Intracystic fluid secretion via ion transporters and channels plays a crucial role in cyst expansion in ADPKD. Unexpectedly, we observed significant and selective up-regulation of NHA2, a member of the *SLC9B* family of Na^+^/H^+^ exchangers that correlated with cyst size and disease severity in ADPKD patients. Using three-dimensional cultures of MDCK cells to model cystogenesis *in vitro*, we show that ectopic expression of NHA2 is causal to increased cyst size. Induction of PC1 in MDCK cells inhibited NHA2 expression with concordant inhibition of Ca^2+^ influx through store-dependent and independent pathways, whereas reciprocal activation of Ca^2+^ influx by a dominant negative, membrane-anchored C-terminal tail fragment of PC1 elevated NHA2. We show that NHA2 is a target of Ca^2+^/NFAT signaling and is transcriptionally induced by methylxanthine drugs such as caffeine and theophylline, which are contraindicated in ADPKD patients. Finally, we observe robust induction of NHA2 by vasopressin, which is physiologically consistent with increased levels of circulating vasopressin and up-regulation of vasopressin V2 receptors in ADPKD. Our findings have mechanistic implications on the emerging use of vasopressin V2 receptor antagonists such as tolvaptan as safe and effective therapy for PKD and reveal a potential new regulator of transepithelial salt and water transport in the kidney.

## Introduction

Autosomal dominant polycystic kidney disease (ADPKD) is a highly prevalent hereditary disease, affecting one in 400–1000 humans (1,2). ADPKD accounts for up to 10% of end-stage renal disease cases, making it one of the leading causes of kidney failure (1). A better understanding of the underlying pathophysiology is key to management of ADPKD and related disorders. ADPKD is caused by mutations in PKD1 and PKD2 encoding polycystin-1 (PC1) and polycystin-2 (PC2), respectively. PC1 is a transmembrane protein with a large extra-cytoplasmic N-terminal domain that is thought to function as a receptor, and a C-terminal cytoplasmic tail (3,4). PC2 is a transmembrane protein that functions as a non-selective Ca^2+^ channel. PC1 interacts with PC2 via C-terminal coil-coiled domains that are required for proper function and trafficking (5,6). The C-terminal cytoplasmic tail of PC1 can undergo proteolytic cleavage and nuclear translocation leading to activation of STAT signaling (7). In renal tubular cells, the polycystins localize to the primary cilium where they regulate intracellular Ca^2+^ and cAMP levels in response to mechano-stimulating urinary flow. A disruption of PC1-PC2 interaction is thought to lead to cyst formation due to abnormal cellular proliferation and fluid secretion, although the specific molecular pathways that connect the underlying genetic defects to disease pathogenesis and cyst development are poorly understood (1,2).

In this context, increased plasma membrane Na^+^/H^+^ exchange (NHE) activity has been reported in cilium-deficient collecting duct cells from the Oak Ridge mouse model of polycystic kidney disease (8,9). Apical mislocalization of NHE1 in these cells leads to hyperabsorption of Na^+^ and epidermal growth factor-induced cell proliferation (8,10). The potential role of other Na^+^/H^+^ exchanger isoforms in PKD pathophysiology has not been explored. Na^+^/H^+^ exchangers belong to the superfamily of monovalent cation/proton antiporters (CPA) that share a common transmembrane organization of 12-14 hydropathic helices with detectable sequence similarity. The CPA1 subgroup includes the electroneutral NHE family of Na^+^(K^+^)/H^+^ exchangers represented by the well-characterized plasma membrane transporter NHE1 (11-13). More recently, a new clade of CPA2 genes was discovered in animals, sharing ancestry with electrogenic bacterial NapA and NhaA antiporters (14,15). Of the two human CPA2 genes, NHA2 has ubiquitous tissue distribution, whereas the closely related NHA1 isoform is restricted to testis (14-16). Despite the multiplicity of genes encoding Na^+^/H^+^ exchangers in mammals, emerging evidence indicates that CPA1 and CPA2 subtypes have distinct, non-redundant transport roles based on differences in chemiosmotic coupling, inhibitor sensitivity and pH activation (17). At the plasma membrane, NHE1 couples proton extrusion to the inwardly directed Na^+^ electrochemical gradient, established by the Na/K-ATPase, to mediate salt reabsorption in epithelia and recovery from acid load. Consistent with these functions, NHE isoforms are activated by low cytoplasmic pH. In contrast, NHA2 has an alkaline pH optimum and appears to couple inward transport of protons to mediate sodium (or lithium) efflux at the cell membrane, recapitulating the function of the bacterial CPA2 ortholog NhaA (12,15,17,18).

Human NHA2 has been implicated as a marker of essential hypertension, with potential roles in kidney diseases relating to salt and water balance (14,18,19). NHA2 expression in kidney is confined to the distal nephron and renal collecting duct, regions that play critical roles in salt, pH and volume homeostasis (15). NHA2 is localized to both principal cells and intercalated cells of the distal tubule *in vivo.* Principal cells maintain sodium and water balance and the intercalated cells control acid-base homeostasis (19). Although the physiological function of NHA2 is largely obscure, defining the pathways regulating NHA2 expression is an important first step towards deciphering its potential role in hypertension and kidney disease.

A key aspect of precision medicine is to harness patient databases to uncover novel risk factors and potential therapeutic targets. This approach led to our unexpected discovery of selective and significant up-regulation of NHA2 expression in polycystic kidney disease. Using an established MDCK model of *in vitro* cystogenesis we show that NHA2 expression correlates with cyst size, replicating patient data. PC1-mediated Ca^2+^ transients and NFAT signaling regulate NHA2 expression, providing a mechanistic link to PKD pathogenesis. To extend physiological relevance, we demonstrate robust vasopressin-mediated NHA2 expression, which is consistent with increased levels of circulating vasopressin and up-regulation of vasopressin V2 receptors in PKD. Importantly, pharmacological inhibition of the NHA2 transporter drastically attenuated cyst size. Taken together, our findings reveal a hitherto under-recognized role for Na^+^/H^+^ exchange activity in PKD progression and allow us to propose NHA2 as a potential chemotherapeutic target.

## Results

### NHA2 is up-regulated in polycystic kidney disease and promotes cyst development *in vitro*

We sought to investigate differences in transcript levels of plasma membrane isoforms of the *SLC9* superfamily of Na^+^/H^+^ exchangers in cysts obtained from patients with autosomal dominant polycystic kidney disease (20). Transcripts of well-studied *SLC9A* (NHE) subfamily genes remained unchanged (NHE1 and NHE3) or showed modest repression (NHE2) (Fig. S1A-C). Unexpectedly, we observed significant up-regulation of a member of the *SLC9B* subfamily, NHA2, relative to normal kidney tissue that correlated with cyst size and disease severity (Fig. 1A): 4.6-fold in large cysts (p=0.002), 3.4-fold in medium cysts (p=0.001), 3.1-fold in small cysts (p=0.0004), and 1.2-fold in minimally cystic tissue (not significant, p=0.49). Based on this observation, we hypothesized that increased expression of NHA2 could contribute to pathophysiology of PKD.

**Fig. 1:**
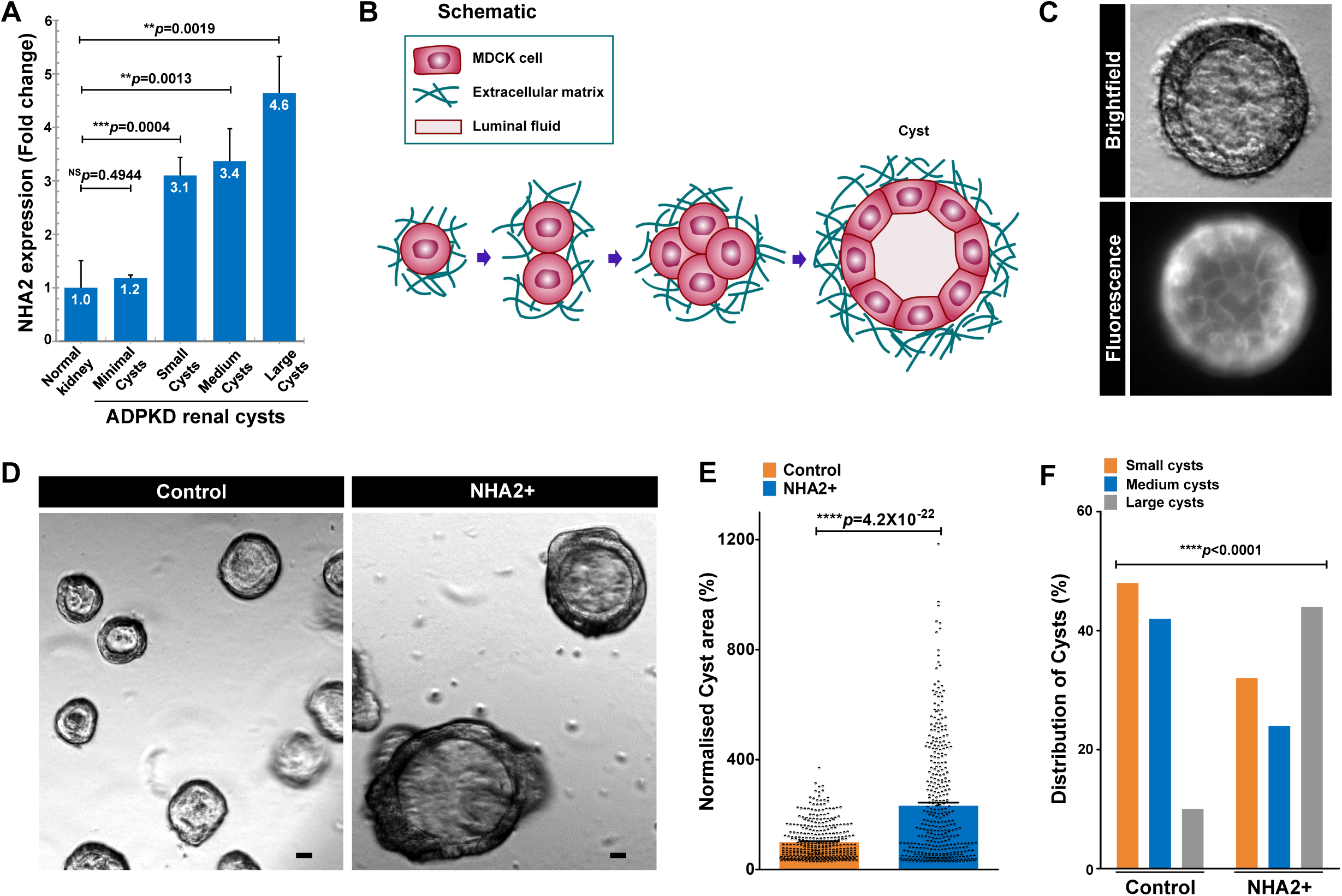
NHA2 is up-regulated in polycystic kidney disease and promotes cyst development *in vitro*. A. Expression profiling of ADPKD patient cysts revealed up-regulation of NHA2 expression relative to normal kidney tissue that correlated with cyst size and disease severity. B. Schematic depicting *in vitro* MDCK cell model of cystogenesis in extracellular matrix (Matrigel). C. Representative image of a MDCK cyst with polarized, single-layer, thinned epithelium surrounding a fluid-filled lumen (*top*; brightfield) with surface expression of NHA2-GFP (*bottom*; fluorescent). D. NHA2+ MDCK cells formed larger cysts (*right*) relative to control (*left*). E. Quantification of cyst size showed significant expansion in NHA2+ cells relative to control on day 10 of culture (Control: 100±66.4; *n*=363, NHA2+: 232.3±213.7; *n*=367, Student’s t test; ^****^*p*=4.2×10^−22^). F. Percentage of large cysts is significantly higher in NHA2+ cells as compared to control. Classification of cysts in three subclasses according to their relative diameter (small/medium/large) showed significantly elevated percentage of cysts with large size in NHA2+ and reduced percentage of cysts with small and medium sizes (χ^2^ test, ^****^p<0.0001).

To test this hypothesis, we used an established *in vitro* MDCK cell model of cystogenesis (21,22). Three-dimensional culturing of MDCK cells in Matrigel produces cysts with polarized, single-layer, thinned epithelium surrounding a fluid-filled lumen (Fig. 1B). Similar to PKD kidneys, MDCK cells in cysts undergo proliferation, fluid transport, and matrix remodeling. To model NHA2 up-regulation, we used MDCK cells stably expressing GFP-tagged NHA2 which colocalizes with the basolateral membrane marker E-cadherin(17) (Pearson’s correlation=0.66±0.11; Manders’ coefficient=0.64±0.17; *n*=25) (Fig. S1D-E). Three-dimensional culturing of these cells in Matrigel resulted in cyst formation with surface expression of NHA2-GFP (Fig. 1C), similar to monolayer cultures. Consistent with our hypothesis linking NHA2 up-regulation to cyst development, we observed significant expansion in cyst size with NHA2 expression relative to non-transfected controls (Fig. 1D-E). Notably, we documented a dramatic difference in the relative cyst diameter on day 10 of culture (Control: 100±66.4; *n*=363, NHA2 up-regulation: 232.3±213.7; *n*=367, *p*=4.2X10^−22^). Intriguingly, we also observed several mutilayered and multiloculated cysts in NHA2+ MDCK cells that were not seen in cysts derived from control MDCK cells (Fig. S1F-G). We classified cysts derived from control cells based on their relative diameter: small (<50 percentile), medium (50-90 percentile), and large (>90 percentile). We observed a significantly higher percentage of larger cysts in NHA2+ cells (control: 10% vs. NHA2 up-regulation: 44%; p<0.0001; Chi-square test; Fig. 1F). Taken together, these findings suggest that NHA2 up-regulation in polycystic kidney disease may promote cyst development.

### Polycystin 1 downregulates Ca^2+^ influx and NHA2 expression

Previously, using a well-characterized MDCK cell model of inducible PC1 expression, we showed that PC1 negatively regulates PC2 levels post-translationally, by enhancing its degradation via the aggresome-autophagosome pathway (23). Using this model, we sought to determine if PC1 induction also alters expression of NHA2 (Fig. 2A). Tetracycline-inducible PC1 expression resulted in significant down-regulation of NHA2 protein levels by ~2.9-fold, relative to non-induced control (p=0.0353; Fig. 2B-C), similar to our previous findings with PC2 (23). Whereas transcript levels of PC2 remained unchanged upon PC1 induction (p=0.663; Fig. 2D), we observed significantly lower NHA2 mRNA levels, consistent with PC1-mediated transcriptional down-regulation of NHA2 (p=0.0029; Fig. 2D). Given that PC2 is a known calcium channel, we next determined if calcium influx is altered by PC1 induction in MDCK cells. We tested two modes of Ca^2+^ entry: store independent Ca^2+^ entry (SICE) and store operated Ca^2+^ entry (SOCE), both mechanisms that are implicated in normal physiology and pathological conditions (24,25). In the absence of PC1 induction, MDCK cells showed robust SICE that was significantly and proportionately attenuated upon induction of PC1 for 24 and 48 hours (Fig. 2E-F). Similarly, PC1 induction for 48 hours also reduced SOCE following thapsigargin-mediated release of store Ca^2+^ (Fig. 2G), consistent with previous reports of PC1 mediated inhibition of STIM1 translocation during store depletion (26). Given these concordant observations on NHA2 expression and Ca^2+^ influx, we investigated whether reduced Ca^2+^ influx in PC1 induced cells is causal to transcriptional down-regulation of NHA2. We first analyzed a publicly available microarray dataset from Ca^2+^-induced activation of primary T lymphoblasts following treatment with Ca^2+^ ionophore ionomycin and phorbol myristate (27). Intriguingly, we observed *SLC9* gene expression changes similar to ADPKD cysts, including ~2.4-fold increase in NHA2 levels, no change in NHE1, and lower, but non-significant levels of NHE2 in response to Ca^2+^ influx in these cells (Fig. S2A-C). Ionomycin treatment increased cytoplasmic Ca^2+^ in PC1 induced MDCK cells (Fig. 2H), resulting in significant and dose dependent increase in NHA2 expression to levels similar to MDCK cells without PC1 induction (Fig. 2I). Taken together these findings are consistent with Ca^2+^-mediated regulation of NHA2 expression by PC1.

**Fig. 2:**
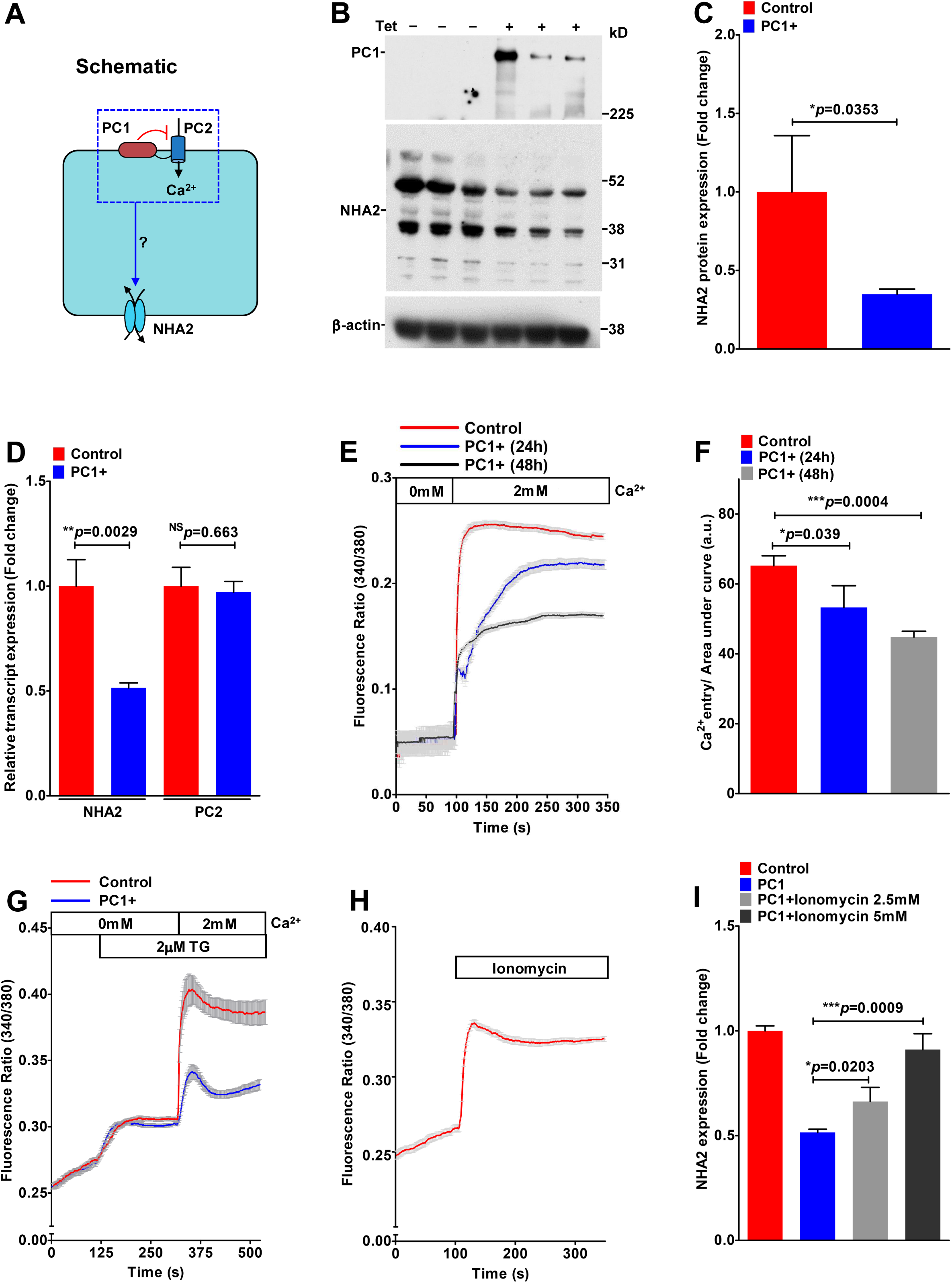
Polycystin 1 downregulates Ca^2+^ influx and NHA2 expression. A. Hypothesis for PC1 mediated inhibition of Ca^2+^ influx through PC2 and NHA2 expression in MDCK cells. B. Western blotting of MDCK cell lysates in the absence or presence of Tet-induced PC1 expression in biological triplicate, probed with antibody to PC1 (*top*), NHA2 (*middle*) and β-actin (*bottom*). (C) Densitometric quantification of Western blot shown in B. NHA2 protein expression was normalized to actin controls and was significantly down-regulated by ~2.9-fold (n=3; Student’s t test; ^*^p=0.0353) in MDCK cells with PC1 induction. D. Quantitative PCR (qPCR) showing significant down-regulation of NHA2 mRNA (n=3; Student’s t test; ^**^p=0.0029) and no change in PC2 mRNA (n=3; Student’s t test; p=0.663) with PC1 induction. (E-F). Representative Fura-2 fluorescence ratio traces (E) and quantitation (F) showing significant and proportionate reduction in store independent calcium entry (SICE) upon induction of PC1 for 24h (n=3; Student’s t test; ^*^p=0.039) and 48h (n=3; Student’s t test; ^***^p=0.0004) in MDCK cells. G. Representative Fura-2 fluorescence ratio traces showing reduction in store operated calcium entry (SOCE) following thapsigargin-mediated release of store Ca^2+^ in MDCK cells with PC1 induction for 48 hours. H. Representative Fura-2 fluorescence ratio traces showing efficacy of Ca^2+^ ionophore ionomycin to enhance cytosolic calcium Ca^2+^ levels. I. qPCR showing significant and dose dependent increase in NHA2 expression with ionomycin treatment to levels similar to MDCK cells without PC1 induction.

### NFAT mediates Ca^2+^-dependent NHA2 expression

The NFAT family of transcription factors has been implicated in the regulation of genes that control numerous aspects of normal physiology including cell cycle progression and differentiation, and in pathological conditions such as inflammation and tumorigenesis (28). The TNF family member <u>R</u>eceptor <u>A</u>ctivator of <u>N</u>F-<u>k</u>B <u>L</u>igand (RANKL) elicits Ca^2+^ oscillations in differentiating osteoclasts that converge on the calcineurin/NFAT pathway(29). During RANKL-induced osteoclast differentiation, NHA2 ranked among top five NFATc1-dependent transcripts, although the physiological role of this up-regulation is yet to be determined (30). To confirm and extend these findings, we analyzed the microarray data from this study (30) and another study reporting NFAT-mediated osteoclast differentiation *in vitro* (31) to show profound, >95% down-regulation of NHA2 levels upon NFAT deletion (Fig. 3A) and in contrast, time-dependent up-regulation of NHA2 levels (~50-fold on day 2-3) upon NFAT activation (Fig. S3A). There was no change in expression of the NHE1 isoform (Fig. 3A). Expression patterns of well-known NFAT target genes, CLCN7 and MMP9 (32), are shown for comparison (Fig. 3A and S3B-C). Consistent with these results, an independent study revealed strong down-regulation of NFATc1 expression and consequently, >30-fold lower NHA2 levels in macrophage colony stimulation factor (M-CSF) and RANKL primed osteoclast precursor myeloid blasts seeded on plastic, relative to cells seeded on bone (33). Similarly, independent studies have reported NFAT-mediated regulation of NHA2 expression in T cells (34,35). Furthermore, NFAT activation might also underlie recent reports of TNF-alpha induced up-regulation of NHA2 expression in human endothelial cells (36).

**Fig. 3:**
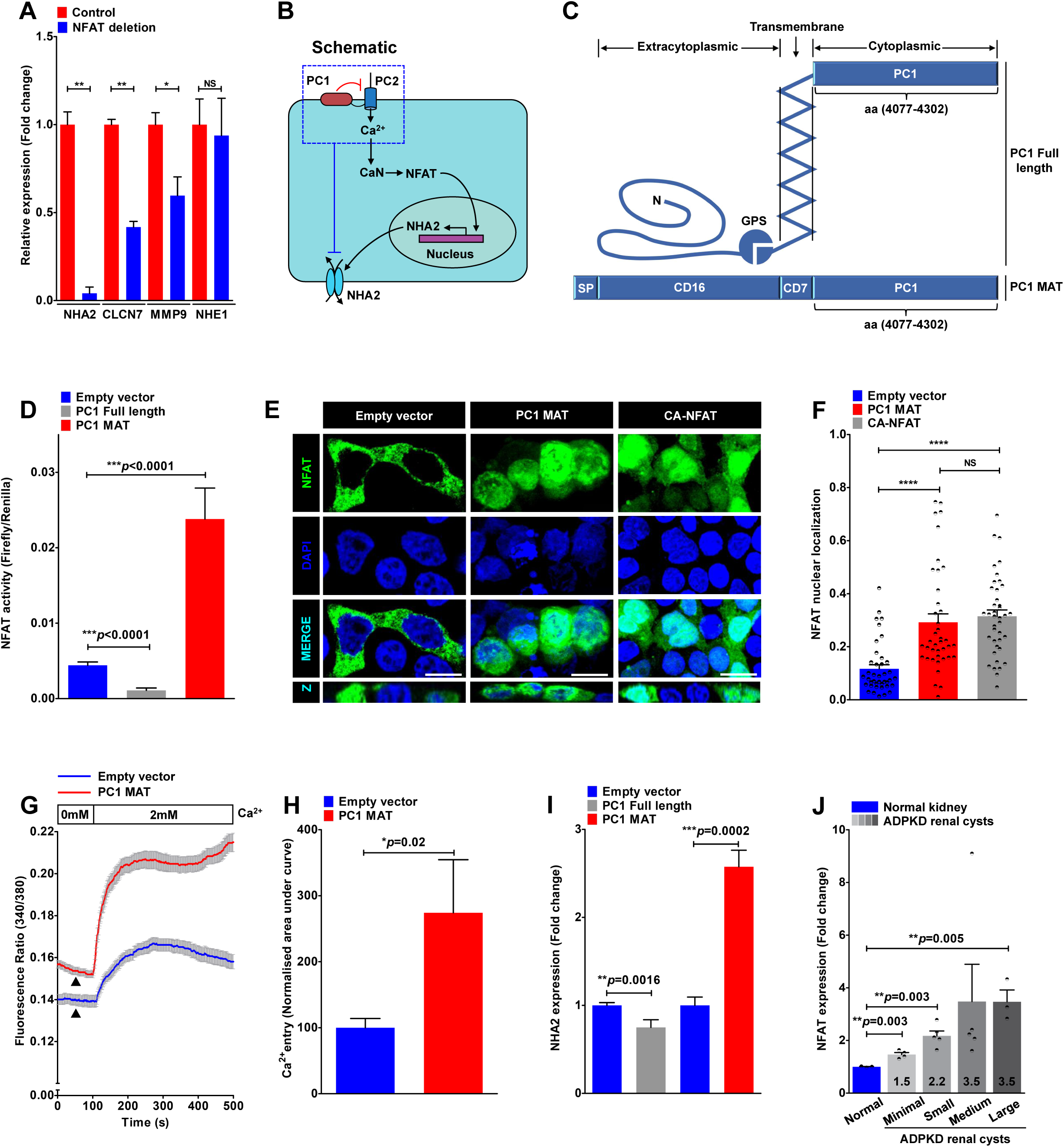
NFAT mediates Ca^2+^-dependent NHA2 expression. A. NFATc1 deletion resulted in profound, >95% down-regulation of NHA2 expression (^**^p=0.0036; Student’s t test) and no change in related NHE1 isoform (p=0.766; Student’s t test) during osteoclast differentiation *in vitro.* Down-regulation of well-known NFAT target genes, CLCN7 and MMP9, are shown for comparison. B. Proposed model for PC1 mediated modulation of Ca^2+^ homeostasis and regulation of NHA2 expression through calcineurin (CaN) and NFAT signaling. C. Schematic of full-length PC1 (*top*) and membrane anchored C-terminal tail fragment of PC1 (PC1-MAT) (*bottom*) containing the C-terminal tail fragment of PC1 linked to the signal peptide (SP) of CD16 and transmembrane domain of CD7. D. Luciferase assay in HEK293 cells to determine NFAT activation using a 3x NFAT binding sequence that drives a firefly luciferase reporter gene and is measured luminometrically. Renilla luciferase driven by a constitutively active SV40 promoter is used to normalize for variations in both cell number and the transfection efficiency. Expression of full-length PC1 resulted in reduction (~4-fold lower; ^***^p<0.0001; n=3; Student’s t test) and of PC1-MAT resulted in increase (~5-fold higher; ^***^p<0.0001; n=3; Student’s t test) in NFAT reporter activity, relative to empty vector transfection control. (E-F). Representative micrographs (E) and quantification using Manders’ coefficient (F) determining fractional colocalization of NFAT-GFP (green) with DAPI (blue). Colocalization is evident in the merge and orthogonal slices (Z) as cyan puncta. In vector transfected cells, NFAT-GFP is predominantly localized in the cytoplasm. Note prominent overlap between NFAT-GFP and DAPI, consistent with increased nuclear translocation, in cells expressing PC1-MAT, similar to cells expressing constitutively active NFAT-GFP (CA-NFAT) (^****^p<0.0001; n=40/each condition; Student’s t test). (G-H). Representative Fura-2 fluorescence ratio traces (G) and quantitation (H) showing ~2.4-fold increase in store independent calcium entry (SICE) in cells transfected with PC1-MAT (n=3; Student’s t test; ^*^p=0.02), relative to empty vector transfection. Note a higher baseline with PC1-MAT, relative to the empty vector control, suggesting higher basal Ca^2+^ levels with PC1-MAT expression. I. qPCR showing ~2.6-fold higher NHA2 expression in cells expressing PC1-MAT (n=3; Student’s t test; ^***^p=0.0002) and ~25%-lower NHA2 levels in cells expressing full-length PC1 (n=3; Student’s t test; ^**^p=0.0016), relative to empty vector transfection. J. NFATc1 expression profiling of ADPKD patient cysts of different sizes revealed up-regulation of NFAT expression relative to normal kidney tissue that correlated with cyst size and disease severity.

Studies have established the importance of NFAT signaling in normal kidney development and in pathological conditions including polycystic kidney disease(37-39). Based on findings from the literature and our data so far, we propose a model that PC1 mediated modulation of Ca^2+^ homeostasis regulates NHA2 expression through NFAT signaling (Fig. 3B). To test this hypothesis, we used a <u>m</u>embrane <u>a</u>nchored C-terminal <u>t</u>ail fragment of PC1 (PC1-MAT) that was previously shown to function as a dominant negative effector and mimic cellular pathologies associated with patient mutations in PC1, including activation of downstream AP-1, WNT, STAT3 and NFAT signaling (7,39-41) (Fig. 3C). We therefore asked if exogenous expression of PC1-MAT fragment could induce NHA2 expression in HEK293 cells. Using the firefly and Renilla luciferase reporter gene system, we first confirmed that ectopic expression of PC1-MAT resulted in significant, ~5-fold increase in NFAT reporter activity, relative to empty vector transfection control (p<0.0001; Fig. 3D). These results were independently validated by visualizing localization of GFP-tagged NFAT: in vector transfected cells, NFAT-GFP is predominantly localized in the cytoplasm, whereas increased nuclear translocation of NFAT-GFP was documented in cells expressing PC1-MAT, similar to cells expressing constitutively active NFAT-GFP (CA-NFAT) (Fig. 3E-F). In striking contrast, expression of full-length PC1 showed significant, ~4-fold down-regulation of NFAT reporter activity, relative to empty vector transfection control (p<0.0001; Fig. 3D). These results are consistent with down-regulation of Ca^2+^ influx and NHA2 expression by induction of full length PC1 in MDCK cells (Fig. 2B-C and 2E-F). We have previously shown that store independent Ca^2+^ entry (SICE) regulates NFAT nuclear translocation and promotes cell proliferation (25). We therefore analyzed SICE in cells transfected with PC1-MAT. In contrast to our SICE data in MDCK cells expressing full-length PC1 (Fig. 2E-F), we observed robust ~2.4-fold increase in Ca^2+^ entry, compared to empty vector control (p=0.02; Fig. 3G-H). We also observed a higher baseline with PC1-MAT, relative to the empty vector control, suggesting higher basal Ca^2+^ levels with PC1-MAT expression (Fig. 3G). Taken together, these data provide strong evidence for reciprocal down- and up-regulation of SICE/NFAT signaling by full length PC1 and PC1-MAT, respectively. Importantly, consistent with our proposed model, we observed ~2.6-fold higher NHA2 expression in cells expressing PC1-MAT (p=0.0002; Fig. 3I), and on the other hand, ~25%-lower NHA2 levels in cells expressing full-length PC1 (p=0.0016; Fig. 3I), relative to empty vector transfection. These findings are also consistent with our data from MDCK cells expressing full-length PC1 (Fig. 2B-D), and evidence from the literature showing significant up-regulation of NHA2 upon ectopic expression of engineered constitutively active NFAT1 in T cells (34). Finally, analysis of NFATc1 expression in ADPKD patient-derived cysts revealed significant up-regulation relative to normal kidney tissue that correlated with cyst size and disease severity (Fig. 3J): large cysts (3.5-fold, p=0.005), medium cysts (3.5-fold, p=0.235), small cysts (2.2-fold, p=0.003), and minimally cystic tissue (1.5-fold, p=0.003). These findings, taken together, suggest that polycystin 1 mediated modulation of Ca^2+^ influx regulates NHA2 expression through the calcineurin/NFAT pathway.

### Potential modulation of kidney function by drug and hormonal regulation of NHA2

Public datasets can be leveraged to identify novel mechanisms of gene regulation, as described earlier (42). To gain new insights into the regulation of NHA2 expression and to predict a functional role of NHA2 in PKD, we performed an unbiased bioinformatics approach to analyze drug-induced gene expression signatures from 1078 microarray studies in human cells. We identified experimental conditions eliciting a minimum of ±2-fold change in NHA2 gene expression (Fig. 4A). Notably, the highest up-regulation of NHA2 (≥3-fold) was observed in response to methyl xanthine drugs: caffeine (7.5mM; 24h) and theophylline (10mM; 24h). Further analysis of these individual microarray studies showed that the ability of these drugs to induce NHA2 expression is dependent on dosage and duration of treatment (Fig. 4B-C). Interestingly, these methyl xanthine drugs are suggested to promote cyst development in PKD patients(43). Furthermore, both caffeine and theophylline are known to activate ryanodine sensitive receptors (RyR) and release Ca^2+^ stores in response to an initial Ca^2+^ entry through PC2 and other plasma membrane Ca^2+^ channels, thereby effectively amplifying cytoplasmic Ca^2+^ (44).

**Fig. 4:**
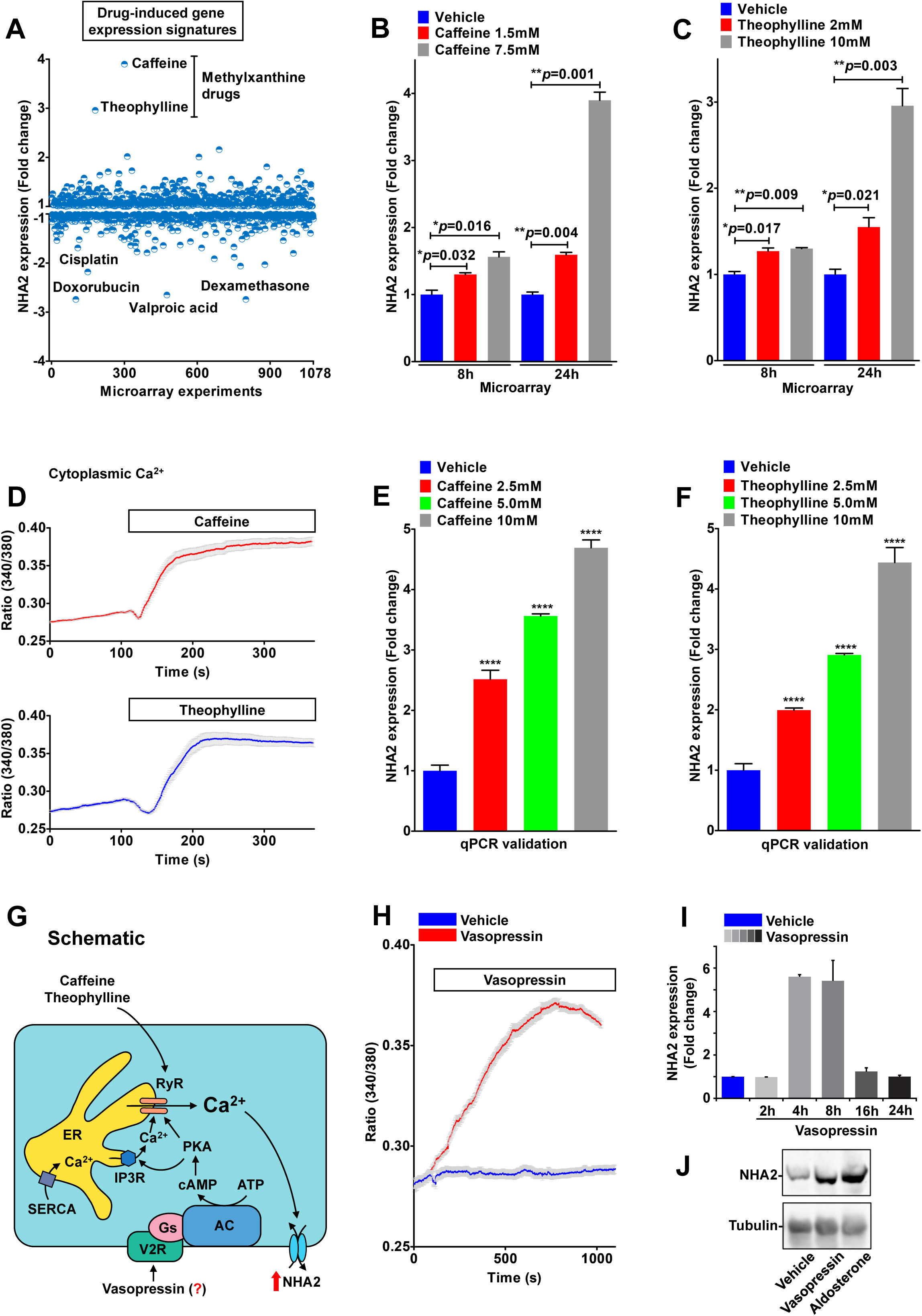
Drug and hormonal regulation of NHA2. A. Expression profiling of NHA2 expression (y axis) obtained from an unbiased bioinformatics analysis of 1078 microarray studies (x axis), as described under “Experimental procedures.” Note that highest up-regulation of NHA2 (≥3-fold) was observed in response to methyl xanthine drugs: caffeine (7.5mM; 24h) and theophylline (10mM; 24h). Four drugs that resulted in maximal down-regulation of NHA2 are valproic acid, dexamethasone, cisplatin and doxorubicin. (B-C). Bar-graphs of NHA2 expression derived from microarray analysis of caffeine (1.5 and 7.5 mM; B) and theophylline (2 and 10mM; C) treatments showing dose and duration (8h and 24h) dependent effects. D. Representative Fura-2 fluorescence ratio traces showing significant increase in cytoplasmic Ca^2+^ with caffeine (*top*) and theophylline (*bottom*) treatment in MDCK cells. (E-F). qPCR analysis to validate bioinformatic studies documenting significant, dose-dependent increase in NHA2 expression with caffeine (E) and theophylline (F), relative to vehicle controls in HEK293 cells. G. Hypothesis for vasopressin mediated regulation of NHA2 expression in renal epithelial cells. Vasopressin stimulation of the V2R receptors results in accumulation of cAMP and activation of PKA. Like caffeine and theophylline, vasopressin-driven PKA activation stimulates Ca^2+^ release via the ryanodine receptor channel from the endoplasmic reticulum (ER). Increased cytosolic Ca^2+^ would in turn increase NHA2 expression. H. Representative Fura-2 fluorescence ratio traces showing significant increase in cytoplasmic Ca^2+^ with vasopressin treatment in MDCK cells, relative to vehicle control. I. qPCR data showing fold change in NHA2 transcript levels following treatment with vasopressin for indicated time periods ranging from 2-24h. Note significant and phasic increase in NHA2 expression with vasopressin treatment that peaked (~6-fold) at 4-8h and reached baseline by 24h. J. Western blot showing NHA2 expression levels following a 6h treatment of MDCK cells with vehicle (*lane* 1), vasopressin (*lane 2*) or aldosterone (*lane* 3), with tubulin serving as a loading control (*bottom panel*).

On the other hand, at least two drugs that resulted in maximal down-regulation of NHA2, namely valproic acid and dexamethasone, are known to reduce cytoplasmic Ca^2+^ concentration (45,46). Interestingly, the therapeutic potential of valproic acid has been shown in zebra fish and mouse models of PKD (47). Taken together, top hits in NHA2 gene expression (both up- and down-regulation) funneled into pathways that modulate cytoplasmic Ca^2+^, further bolstering the credibility of our model of Ca^2+^ mediated regulation of NHA2 expression. Our previous studies show that NHA2 is an important cellular lithium efflux transporter that might be involved in Li^+^ clearance in kidney (17). In this context it is important to note that methylxanthine drugs caffeine and theophylline that increase NHA2 levels are well known to promote Li^+^ clearance via kidney. These drugs enhance renal salt and water transport and are contraindicated for Li^+^ therapy (48,49). Using ratiometric Fura 2 Ca^2+^ imaging, we first confirmed significant increase in cytoplasmic Ca^2+^ with caffeine and theophylline treatment in MDCK cells (Fig. 4D). Furthermore, caffeine and theophylline treatment elicit significant, dose-dependent increase in NHA2 expression (Fig. 4E-F), relative to vehicle controls, validating the findings from bioinformatic studies (Fig. 4A-C).

**Fig. 5:**
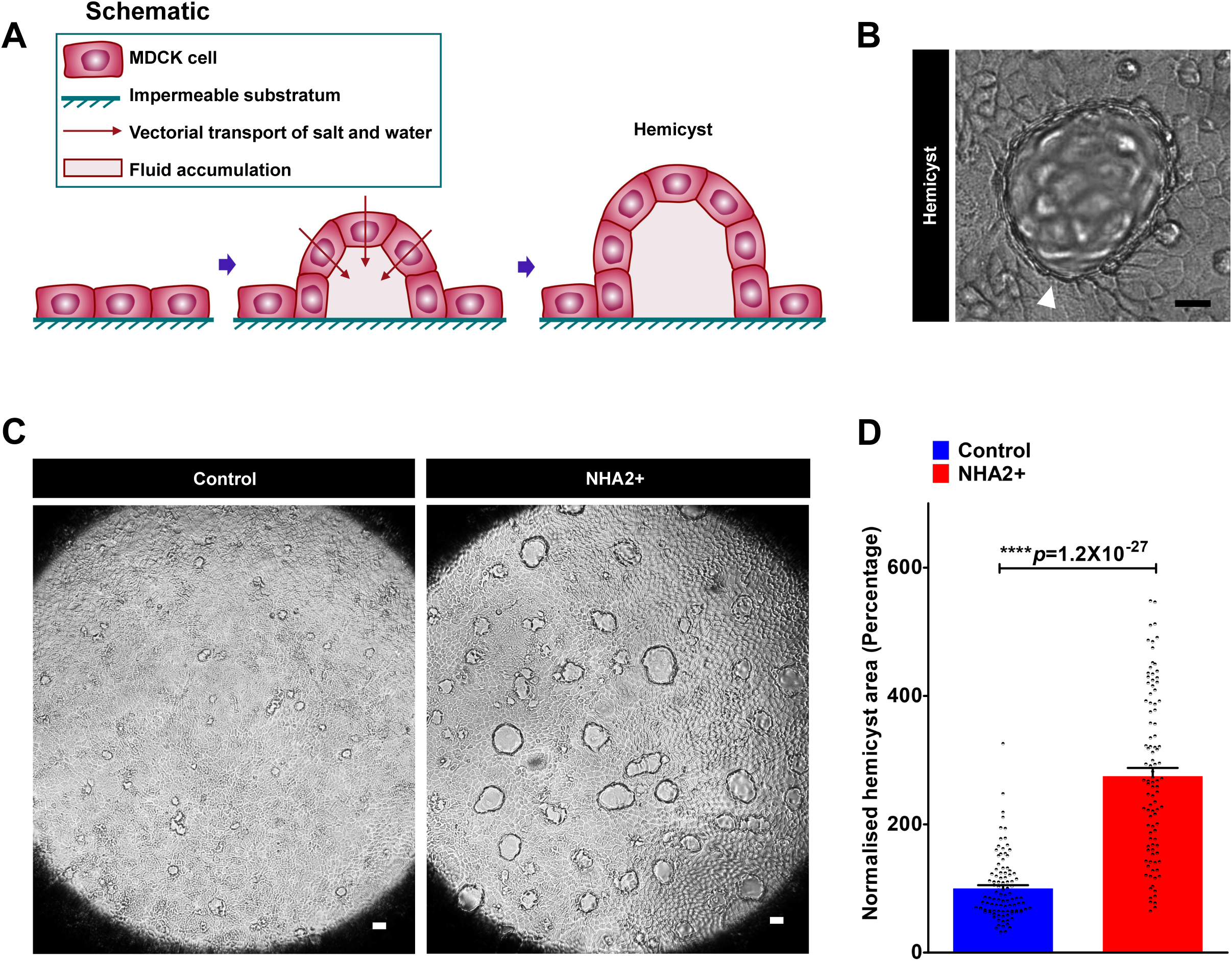
NHA2 promotes hemicyst formation. A. Schematic depicting generation of fluid-filled hemi-cysts or domes with MDCK cells.
B. Representative image of a MDCK hemicyst with fluid accumulation focally beneath the epithelium (*white arrow*). C. NHA2+ MDCK cells formed larger hemicysts (*right*) relative to control (*left*). D. Quantification of hemicyst development showed significant expansion in hemicyst area with NHA2 expression relative to control (Control: 100±5.01; *n*=100, NHA2+: 275.1±12.75; *n*=100, Student’s t test; ^****^p=1.2×10^−27^), suggesting an increase in vectorial transport of salt and water with NHA2 expression.

Given that physiological concentrations of vasopressin, a hormone strongly implicated in PKD pathogenesis and a promising drug target, also functions via activation of RyR (50), we considered whether vasopressin regulates NHA2 expression (Fig. 4G). The MDCK cell line is a well-established model to study transport function changes in the distal renal tubule in response to hormones, including vasopressin and aldosterone (51,52). To test our hypothesis, we used the MDCK cell model and first documented robust increase in cytoplasmic Ca^2+^ following vasopressin treatment, relative to vehicle control (Fig. 4H). Compared with the quick (~50-100s) initial slope of the Ca^2+^ rise in caffeine and theophylline treatment, the slope of the Ca^2+^ rise was more gradual (~500-1000s) consistent with indirect activation of downstream signaling cascades by vasopressin (Fig. 4G). Of note, the peak heights of cytosolic [Ca^2+^] were comparable between caffeine, theophylline and vasopressin. It is worth noting that both caffeine and vasopressin have been shown to activate NFAT nuclear translocation and signaling (53,54). Therefore, to determine if rise in cytosolic Ca^2+^ with vasopressin is translated to increase in NHA2 expression, we treated MDCK cells with vasopressin for different intervals, ranging from 2-24h. We observed significant and phasic increase in NHA2 expression with vasopressin treatment that peaked (~6-fold) at 4-8h and reached baseline by 24h (Fig. 4I). Using Western blotting we confirmed robust elevation of NHA2 protein with vasopressin treatment for 6h (Fig. 4J).

These findings are consistent with our recent *in vivo* studies showing that high salt diet in mice significantly increased transcript and protein expression of NHA2 in renal tubules (19). High salt diet significantly raises circulating levels of vasopressin to retain water and help sustain normal plasma osmolarity (55). Likewise, increased vasopressin concentrations have been associated with PKD severity and disease progression and in experimental studies vasopressin has been shown to directly modulate transepithelial fluid transport and regulate cyst growth (50). Intriguingly, studies have documented that the NHA2 inhibitor phloretin attenuates vasopressin-stimulated solute movement (56). Thus, NHA2 might regulate salt and water transport across the epithelium and help contribute to the functions of vasopressin in PKD and high salt diet, both are associated with hypertension.

For comparison, we tested the effect of another hormone, aldosterone, on NHA2 expression in MDCK cells. Basic and clinical research have shown increased activation of the renin-angiotensin-aldosterone-system in ADPKD (57,58). Intriguingly, like vasopressin, aldosterone also increased NHA2 mRNA (Fig. S4A) and protein levels (Fig. 4J). Similar induction with both vasopressin and aldosterone expression has been reported in the literature for aquaporin-2 and gamma subunit of the epithelial sodium channel (ENaC)(59,60). Moreover, a synergistic effect of vasopressin and aldosterone has been described for Na^+^ channel and Na^+^/K^+^-ATPase activities (61,62). It is worth noting that aldosterone has been shown to enhance global Ca^2+^ transients, through activation of protein kinase A (PKA)(63). To evaluate functional consequences of the hormonal induction of NHA2, we monitored growth-sensitivity to lithium in MDCK cells. Previously, we showed that NHA2 is functionally coupled to plasma membrane V-ATPase in MDCK cells to mediate robust H^+^-driven efflux of Li^+^, resulting in selective survival and growth advantage in media with high concentration of lithium (17). Consistent with increased NHA2 mRNA and protein expression, treatment with either vasopressin or aldosterone for 8 h resulted in increased cell survival in media supplemented with 90 mM LiCl, relative to untreated control (Fig. S4B).

Vasopressin and aldosterone play key roles in the fine adjustment of transport of salt and water in the nephron (60-62). In order to test the potential role of NHA2 in regulating transepithelial fluid transport, we generated fluid-filled hemi-cysts or domes, a simple, yet powerful assay widely used over the last 40 years to determine vectorial salt and water transport *in vitro* (64,65). MDCK cells grown on impermeable substratum, such as plastic, spontaneously form hemicysts due to transepithelial transport and accumulation of fluid in focal regions, beneath the cell layer (Fig. 5A-B). Studies have established that cyclic AMP levels, Na^+^/H^+^ exchange activity and PC1 function regulate hemicyst formation (65-67). Importantly, vasopressin and aldosterone treatment promote transepithelial transport and robustly stimulate hemicyst formation in MDCK cells(68,69). Consistent with these findings, we show that ectopic expression of NHA2 in MDCK cells resulted in larger, ~2.8-fold increase in area of hemicyst (Fig. 5C-D), suggesting an increase in vectorial transport of salt and water with NHA2 expression that might, in part, contribute to the downstream effects of vasopressin and aldosterone in PKD.

### Knockdown or inhibition of NHA2 attenuates cyst development *in vitro*

Given that fluid secretion and cyst expansion are major factors for the progressive and irreversible decline and renal failure in PKD, therapies targeting salt and water transport are of major interest as an alternative, or to complement anti-proliferative therapies in PKD (70). To evaluate the therapeutic potential of targeting NHA2 in PKD we used two complementary approaches: shRNA-mediated knockdown of NHA2 and pharmacological inhibition of NHA2 using phloretin. MDCK cells have low endogenous NHA2 expression (17), therefore we knocked down NHA2 in NHA2+ MDCK cells (Fig. S5A). We have previously shown that overexpression of NHA2 does not alter growth in MDCK cells (17). Consistent with these findings, we now show that lentivirally mediated knockdown of NHA2 also did not alter growth of MDCK cells (Fig. 6A). Next, we tested the effect of NHA2 knock down on cyst formation by MDCK cells (Fig. 6B-C). Specific knockdown of NHA2, using two different lentiviral shRNA constructs, resulted in formation of significantly smaller (~70-80% lower diameter) cysts (Fig. 6B-C).

**Fig. 6:**
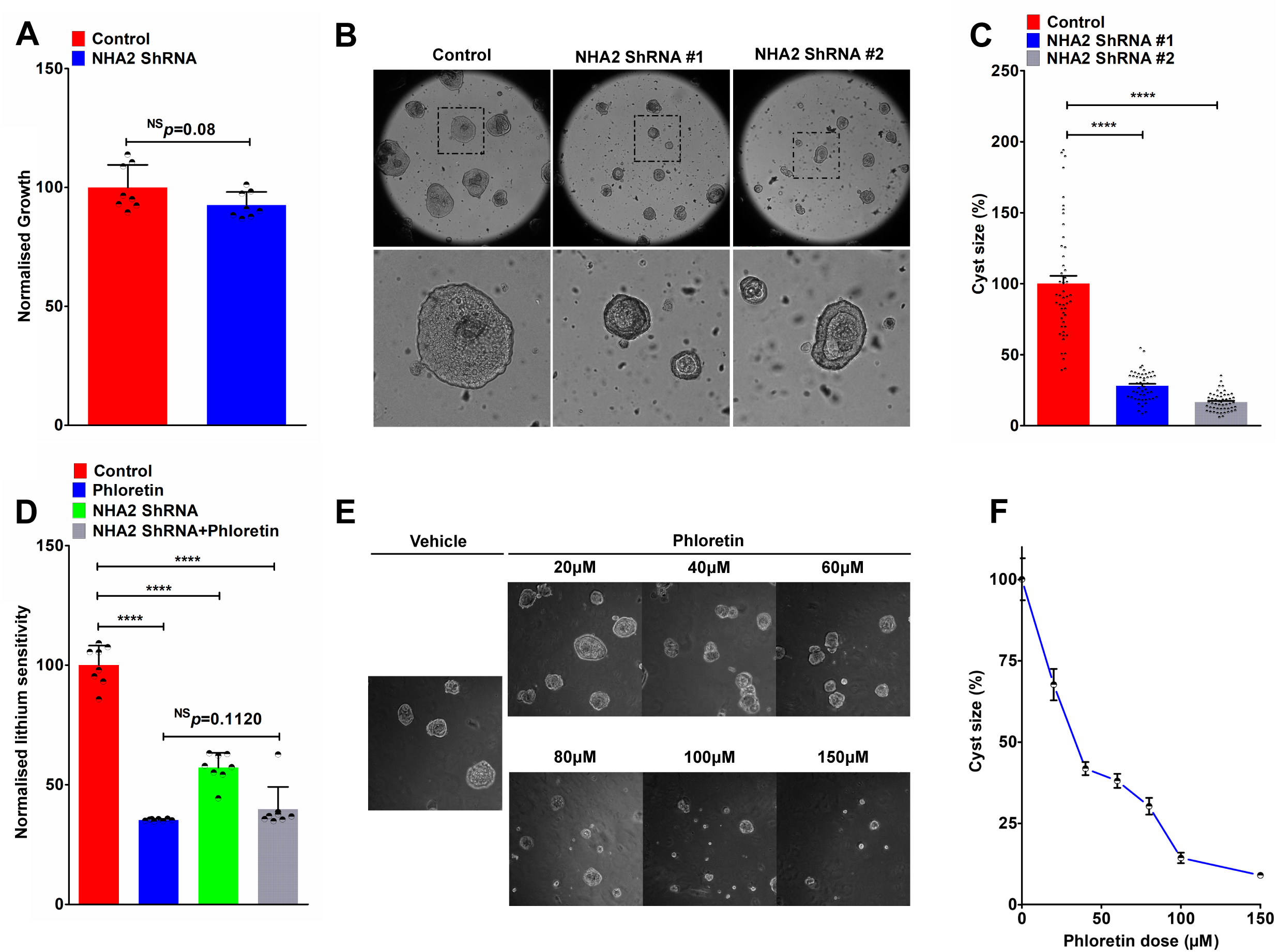
Knockdown or inhibition of NHA2 attenuates cyst development *in vitro*. A. Lentivirally mediated knockdown of NHA2 does not alter growth of MDCK cells. Cell growth activity was measured by MTT assay, normalized to the scrambled control (p=0.08; n=8; Student’s t test). B. Lentivirally mediated specific knockdown of NHA2, using two different lentiviral shRNA constructs (ShRNA#1 and ShRNA#2), resulted in formation of significantly smaller cysts, relative to control. Representative images are shown, with bottom row showing zoom of boxed region from top row. C. Quantification of cyst development showing significant reduction in cyst size with NHA2 depletion relative to control on day 10 of culture (Control: 100.0±5.6; *n*=50, NHA2 ShRNA#1: 28.0±1.46; n=50, NHA2 ShRNA#2: 16.7±0.91; n=50, Student’s t test; ^****^*p*<0.0001). D. Lithium sensitivity assay to validate the effect of phloretin. Cell Survival in the presence of 100 mM LiCl was measured using MTT. Phloretin treatment and lentiviral NHA2 knockdown conferred significant growth sensitivity to lithium (n=8; Student’s t test; ^****^*p*<0.0001). Note no difference in lithium sensitivity between phloretin treatment with or without NHA2 knockdown (n=8; Student’s t test; *p*=0.1120). E. Phloretin (20-150μM) mediated NHA2 inhibition resulted in a dose-dependent reduction in cyst size in Matrigel. Representative images are shown, with the vehicle treatment on the left. F. Quantification of cyst development showed significant reduction in cyst size with phloretin mediated NHA2 inhibition relative to vehicle treatment on day 10 of culture (n=50/each condition; Student’s t test; ^****^*p*<0.0001).

NHA2 is insensitive to classical NHE inhibitors such as amiloride and its derivatives but is sensitive to phloretin (17). In contrast to the effect of pharmacological inhibition of NHE1, inhibition of NHA2 with phloretin did not significantly alter cytoplasmic pH (Fig. S5B-C). Therefore, to functionally validate the effect of phloretin, we turned to lithium sensitivity as a defining phenotype of NHA2 transport function. Treatment of NHA2+ MDCK cells with phloretin conferred growth sensitivity to lithium (Fig. 6D). Notably, no difference in lithium sensitivity was observed between phloretin treatment with or without NHA2 knockdown indicating the specificity of lithium sensitivity assay as a marker of NHA2 activity (Fig. 6D). Next, we analyzed the effect of phloretin (20-150μM) on MDCK cyst formation in Matrigel. Consistent with lower cyst size resulting from NHA2 knockdown (Fig. 6B-C), we observed dose-dependent attenuation of cyst size with NHA2 inhibition with phloretin (Fig. 6E-F). Taken together, these findings suggest that NHA2 may be a potential drug target for attenuating cyst development in PKD.

## Discussion

The three key components of cystogenesis in PKD are abnormalities in tubular cell proliferation, matrix remodeling and fluid secretion (70). Cysts arise from the renal tubule as saccular expansions, and eventually become pinched off and anatomically separated from the tubule. Thus, the intracystic fluid is derived from transepithelial fluid secretion due to abnormal reversal of normally absorptive epithelium to a secretory one. This leads to dramatic expansion of cystic volume, which is the single most important predictor of progressive deterioration and renal failure in PKD (70,71). Therefore, a better understanding of pathways driving salt and water transport in PKD models could lead to novel strategies to halt disease progression. Although Na^+^/H^+^ exchangers are known to be of vital physiological importance for fluid and salt homeostasis in the kidney (11,12), their contribution to the pathology of PKD has yet to be fully explored or recognized. Using *in vitro* models of cystogenesis in conjunction with gene expression data from PKD patients, we now show that NHA2 expression is regulated by PC1 via Ca^2+^ and NFAT signaling to promote cyst development. A better understanding of a kidney-specific role for NHA2 in PKD awaits a mechanistic study of transport. It is as yet unclear whether the normal physiological role of NHA2 in renal tubular cell is in salt and water reabsorption or secretion or both depending on the coupling ion (sodium or protons) and localization (apical or basolateral)(19).

Although far from conclusive, there are tantalizing hints pointing to a role for NHA2 in renal function and hypertension that may be relevant to pathophysiology of PKD. First, *SLC9B2*, the gene encoding NHA2, lies within a limited (~300 genes) region of human chromosome 4q24 genetically linked to phloretin-sensitive and amiloride-insensitive sodium-lithium countertransport (SLC), an activity linked to essential hypertension (14,18). Indeed, NHA2 mediates phloretin-sensitive and amiloride-insensitive SLC activity in stably transfected MDCK cells (17). SLC activity has been associated with PKD and there is an intriguing, yet unexplained link between lithium treatment and formation of renal cysts in human and rodents (72-74). Second, a recent genome-wide association study (GWAS) of renal function identified NHA2 as a genetic risk locus for estimated glomerular filtration rate by serum creatinine. GFR, which shows heritability in the range of 36-75%, is commonly used to monitor kidney function decline in PKD (75). Third, NHA2 expression is predominant in the distal nephron where sodium reabsorption is under hormonal and dietary control and, consistent with the induction of renal NHA2 expression by a high salt diet in mouse (15,19). In addition to these findings, we now show that NHA2 is highly elevated in PKD, a disease, with predisposition to hypertension (57).

The dysregulation of two second-messengers, Ca^2+^ and cAMP, is central to the disease mechanism in PKD and there is evidence connecting NHA2 to both. Here we show that methylxanthine drugs (caffeine and theophylline) that activate Ca^2+^ release via the ryanodine receptor channel (RyR), increase NHA2 expression which is associated with cyst growth (Fig. 7). Furthermore, we show that NHA2 is up-regulated by vasopressin, a hormone that in physiological concentrations functions through activation of RyR and increases cytoplasmic Ca^2+^ levels. Thus, NHA2 is a novel vasopressin effector in renal epithelial cells with potential implications for PKD.

**Fig. 7:**
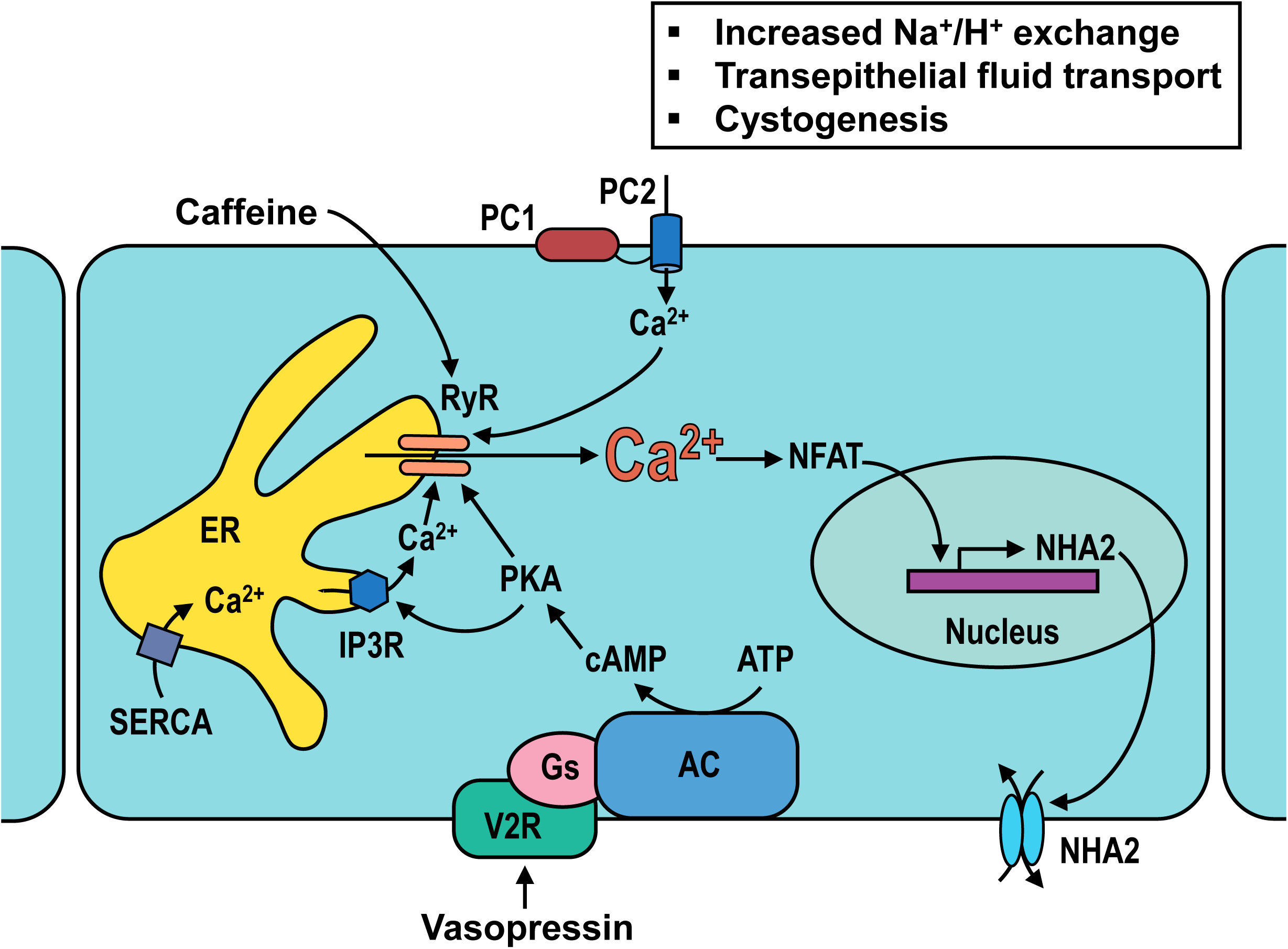
Proposed role for NHA2 in PKD. Polycystins 1 and 2 (PC1 and PC2) regulate Ca^2+^ influx, NFAT signaling and NHA2 expression. Methylxanthine drugs (caffeine and theophylline) activate ryanodine sensitive receptor channel (RyR) and release Ca^2+^ stores from the endoplasmic reticulum (ER) in response to an initial Ca^2+^ entry through PC2 and other plasma membrane Ca^2+^ channels, thereby effectively amplifying cytoplasmic Ca^2+^ and increasing NHA2 expression. Vasopressin stimulation of the V2R receptors results in accumulation of cyclic AMP (cAMP) and activation of protein kinase A (PKA). Like caffeine and theophylline, vasopressin-driven PKA activation stimulates Ca^2+^ release via the RyR, increase NHA2 expression which is associated with cyst growth.

The precise NHA2-mediated regulation of transepithelial fluid transport and cystogenesis remains to be determined. Vasopressin is secreted in response to increased plasma osmolality resulting from high salt intake, consistent with our previous observation that high sodium diet in mice significantly increases expression of NHA2 in renal tubules (19,55). Previous studies have shown that membrane anchored C-terminal tail fragment of polycystin 1 (PC1-MAT) enhances caffeine-induced intracellular Ca^2+^ levels (39). Similarly, caffeine has been reported to synergize the effects of desmopressin, a synthetic analogue of vasopressin (43). Caffeine is also thought to promote cyst development in PKD patients (43). Furthermore, caffeine and theophylline are well known to promote clearance of lithium through kidney (48,49). Given our previous findings that NHA2 is an important cellular lithium efflux transporter (17), we suggest that induction of NHA2 expression might, in part, contribute to the downstream effects of caffeine in PKD and lithium clearance.

NHA2 may mediate cross-talk between cAMP and Ca^2+^ signaling. In spermatozoa, loss of NHA2, or the closely related NHA1 isoform, led to decreased soluble adenylyl cyclase and cAMP levels, and conversely, high levels of NHA antiporters stabilized soluble adenylyl cyclase expression (16). Given the central role of cAMP in cyst formation and growth, elevated NHA2 could exacerbate cAMP signaling in ADPKD patients, which might in turn modulate the activity of other transport proteins. PKD patients have high circulating levels of vasopressin. Treatment with vasopressin V2-receptor antagonist tolvaptan has been shown to inhibit cyst progression in preclinical studies and large clinical trials (50). The potential synergistic or additive effects of NHA2 inhibition with tolvaptan merits further study. In summary, our findings warrant a thorough *in vivo* investigation of NHA2 as a potential new player in cellular salt and pH regulation in the kidney and in the pathogenesis of PKD and hypertension.

## Materials and Methods

### Cell culture

Stable MDCK cell line with inducible PC1 expression was generated using Flp-In System (Invitrogen) by transfecting pcDNA5/FRT/TO-based wild-type mPkd1 expression plasmid according to the manufacturer’s protocol (23). These cells were cultured in DMEM supplemented with 10% fetal bovine serum, 150 μg/ml hygromycin, and 5 μg/ml blasticidin, with PC1 expression induced with tetracycline (2 μg/ml). Stable MDCK cell line with NHA2 expression was generated by transfecting pEGFPC2 vector carrying human NHA2 tagged to GFP at the N terminus and selecting for G418 antibiotic resistant clones (17). MDCK cells (control and NHA2+) were cultured in MEM supplemented with 10% fetal bovine serum. HEK293 cells were cultured in DMEM supplemented with 10% fetal bovine serum. Culture conditions were in a 5% CO2 incubator at 37°C. 3-dimensional cysts were cultured in 10% Matrigel with a 100% Matrigel basement membrane and grown in full media, which was changed every 2 days (21,22). MDCK hemicysts in monolayer cultures were generated as previously described (65,66). Hormonal treatment protocols (vasopressin-1μM or aldosterone-100nM) were previously reported to regulate transporter expression and function in MDCK cells (51,52). Lithium sensitivity was determined by measuring cell growth in the presence of media containing 90-100 mM LiCl. Cell growth was quantified using MTT assay (ThermoFisher Scientific), following the manufacturer’s instructions (17).

### Plasmids and transfection

PC1 full-length plasmid was a gift from Dr Gregory Germino (Addgene plasmid #21369). Membrane anchored C-terminal tail fragment of PC1 (PC1-MAT) plasmid was a gift from Dr Thomas Weimbs (Addgene plasmid #41567). GFP-tagged NFAT1 (1–460) plasmid was described previously(25). Constitutively active NFAT (CA-NFAT) was a gift from Dr Anjana Rao (Addgene plasmid #11102). pGL3-NFAT-luc was a gift from Dr Jerry Crabtree (Addgene plasmid #17870). pSV40-RL was a gift from Dr Jennifer L. Pluznick (Johns Hopkins University). HsNHA2 short hairpin RNA (shRNA) sequence (5’-CACTGTAGGCCTTTGTGTTGTTCAAGAGACAACACAAAGGCCTACAGTTTTTTCAATT-3’) was designed to knockdown NHA2 (ShRNA#1). An additional knockdown construct (ShRNA#2) targeting NHA2 with sequence (5’-CCGGGCATTGCAGTATTGATACGAACTCGAGTTCGTATCAATACTGCAATGCTTTTTTG-3’) was purchased from Sigma (#TRCN0000130075). MDCK cells were transfected using lentiviral packaging and expression. HEK293 cells were transfected using Lipofectamine 2000 reagent, as per the manufacturer’s instructions.

### Antibodies and reagents

Specific rabbit polyclonal NHA2 antibody was raised against a 15-aa peptide of NHA2 (14,17). Mouse monoclonal antibodies used were Anti-PC1 (7E12) (#sc-130554, Santa Cruz Biotechnology), Anti-α-Tubulin (#T9026, Sigma), Anti-β-Actin (#A5441, Sigma), and Anti-e-cadherin (#610181, BD Transduction Laboratories). Aldosterone (A9477), Blasticidin (#15205), Caffeine (#C0750), Ionomycin (#I3909), Lithium chloride (#L9650), Phloretin (#P7912), Thapsigargin (#T9033), Theophylline (#T1633), and Vasopressin (#V9879) were obtained from Sigma. Tetracycline Hydrochloride (A1004-5) and Hygromycin B (#10843555001) were purchased from Zymo Research and Roche, respectively.

### Bioinformatics

Public datasets were mined to uncover novel mechanisms of gene regulation, as described earlier (42). We analyzed drug-induced gene expression signatures from 1078 microarray studies in human cells to identify drugs that significantly altered NHA2 expression. Experimental conditions eliciting a minimum of ±2-fold change in gene expression were selected for further pathway analysis. Normalized gene expression data was obtained from Genevestigator (Nebion AG). Other mammalian gene expression datasets included in the study are GSE7869, GSE37219, GSE50971 and GSE57468.

### Cytoplasmic pH measurement

Cytoplasmic pH was measured using pHrodo Red AM Intracellular pH Indicator (#P35372, Thermo Fisher Scientific), following manufacturer’s instructions. Briefly, cells were rinsed with serum-free medium and then incubated with 1 μL/ml of pHrodo Red AM at 37°C for 30 minutes. Cells were washed, trypsinized and pH was determined by flow cytometry analysis of ~10,000 cells in biological triplicates using the FACSCalibur instrument (BD Biosciences). Cells were gated on a forward scatter and side scatter to obtain live single cells. A four-point calibration curve with different pH values (4.5, 5.5, 6.5 and 7.5) was generated using Intracellular pH Calibration Buffer Kit (#P35379, Thermo Fisher Scientific) in the presence of 10μM K^+^/H^+^ ionophore nigericin and 10μM K^+^ ionophore valinomycin.

### Quantitative Real-time PCR (qPCR)

mRNA was isolated using RNeasy Mini kit (#74104, Qiagen) with DNase1 (#10104159001, Roche) treatment and complementary DNA was synthesized using the high-Capacity RNA-to-cDNA Kit (#4387406, Applied Biosystems), following the manufacturer’s instructions. Gene expression was assessed by quantitative real-time PCR using following Taqman gene expression assays: Dog: NHA2, Cf02729492_m1; PC2, Cf02690612_m1; GAPDH, Cf04419463_gH; Human: NHA2, Hs01104990_m1; GAPDH, Hs02786624_g1.

### Western blot

Cells were harvested and lysed with 1% Nonidet P-40 (Sigma) supplemented with protease inhibitor cocktail (Roche). Following sonication, cell lysates were centrifuged for 15 min at 14,000 rpm at 4°C. The protein concentrations were measured using the BCA assay. Equal amounts of total protein were electrophoretically separated by polyacrylamide gel (NuPAGE) before transferring onto nitrocellulose membranes (Bio-Rad). The membranes were blocked with 5% milk, followed by overnight incubation with primary antibodies and 1h incubation with HRP-conjugated secondary antibodies (GE Healthcare). Amersham 600 chemiluminescence imager system was used to capture images.

### Confocal microscopy

To determine colocalization of NHA2-GFP with E-cadherin, cultured MDCK NHA2+ cells on polyornithine coverslips were pre-extracted with PHEM buffer (60 mM PIPES, 25 mM HEPES, 10 mM EGTA, and 2 mM MgCl2, pH 6.8) containing 0.025% saponin for 2 min, then washed twice for 2 min with PHEM buffer containing 0.025% saponin and 8% sucrose. Following fixation (4% paraformaldehyde and 8% sucrose in PBS) for 30min, cells were blocked with 1% BSA and 0.025% saponin in PBS for 1 hr. Cells were stained with primary Anti-e-cadherin antibody (overnight) and Alexa Fluor 568 conjugated secondary antibody (1 hr). On the other hand, to monitor nuclear translocation of NFAT, cultured HEK293 cells with NFAT-GFP transfection were fixed with a solution of 4% paraformaldehyde in PBS. Following DAPI staining, coverslips from both experiments were mounted onto slides using Dako fluorescent mounting medium and were imaged using LSM 700 Confocal microscope (Zeiss). Confocal imaging was performed with a ×63 oil immersion objective and the fractional colocalization was determined using the JACoP ImageJ plugin.

### Calcium Imaging

Calcium imaging was performed as we previously described (25). Briefly, cells were cultured on 25 mm circular coverslips. Cells were briefly washed with PBS and loaded with Fura-2 AM at 1 mg/ml in calcium imaging buffer (126 mM NaCl, 4.5 mM KCl, 2 mM MgCl_2_, 10 mM Glucose, 20 mM HEPES, pH 7.4) containing 2 mM CaCl_2_. After incubation at room temperature for 30 minutes, cells were washed briefly in imaging buffer without Fura-2 AM. After baseline recordings were established, cells were excited at 340 nm and 380 nm, and Fura-2 AM emission at 505 nm was monitored.

### Luciferase assay

Luciferase assay was performed as we previously described (42). Briefly, pGL3-NFAT-luc (encoding a 3x NFAT binding sequence and firefly luciferase) and pSV40-RL (encoding for a constitutively active SV40 promoter and Renilla luciferase) plasmids were transiently transfected into HEK293 cells using lipofectamine 2000 reagent, as per manufacturer’s instructions. The ratio of firefly luciferase and Renilla luciferase, a measure of NFAT activity, was determined using the Dual Luciferase Assay System (#E1910, Promega). Data were collected using a FLUOstar Omega automated plate reader (BMG LabTech).

## Acknowledgements

We gratefully acknowledge the Baltimore Polycystic Kidney Disease (PKD) Research and Clinical Core Center for stable MDCK cells with inducible PC1 expression. This work was supported by grants from the National Institutes of Health to R.R. (DK108304) and to V.C. (DK103078). H.P. was a Fulbright Fellow supported by the International Fulbright Science and Technology Award. K.C.K. was a postdoctoral fellow of the American Heart Association.

## Figure legends

**Fig. S1:**
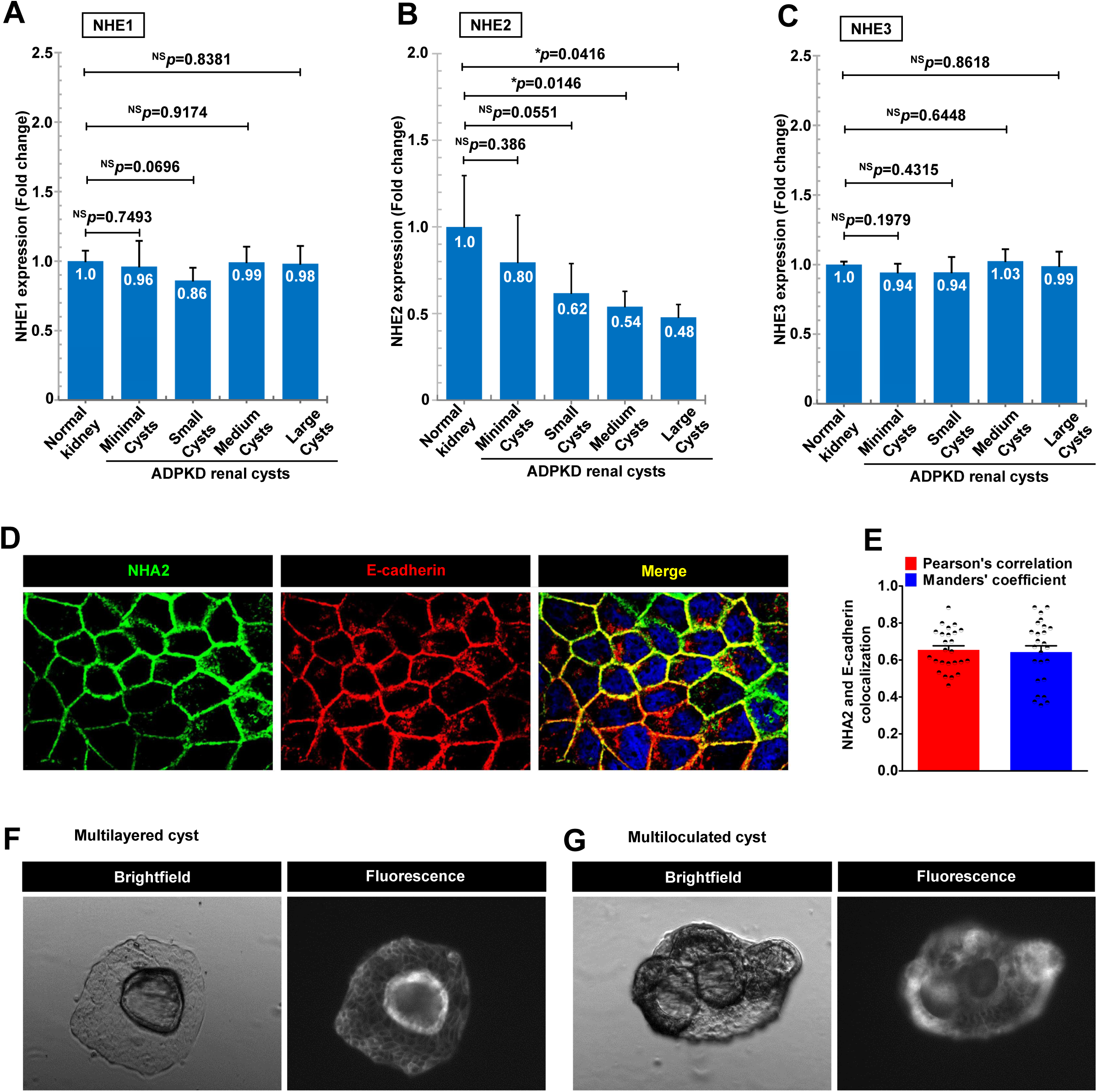
Expression of *SLC9A* (NHE) genes in ADPKD patient cysts. (A-C). Expression profiling of plasma membrane *SLC9A* (NHE) genes in ADPKD patient cysts of different sizes showed no change (NHE1 and NHE3) or showed modest repression (NHE2). (D-E). Representative confocal microscopic images (D) and quantification (E) showing colocalization of NHA2-GFP (*green*) with basolateral marker E-Cadherin (*red*) in DAPI (*blue*) stained cells. Colocalization is evident in the merge as *yellow* puncta (Pearson’s correlation=0.66±0.11; Manders’ coefficient=0.64±0.17; n=25). (F-G). Representative images of MDCK cysts to document several mutilayered and multiloculated cysts in NHA2+ MDCK cells (*left*; brightfield) with surface expression of NHA2-GFP (*right*; fluorescent) that were not seen in cysts derived from control MDCK cells.

**Fig. S2:**
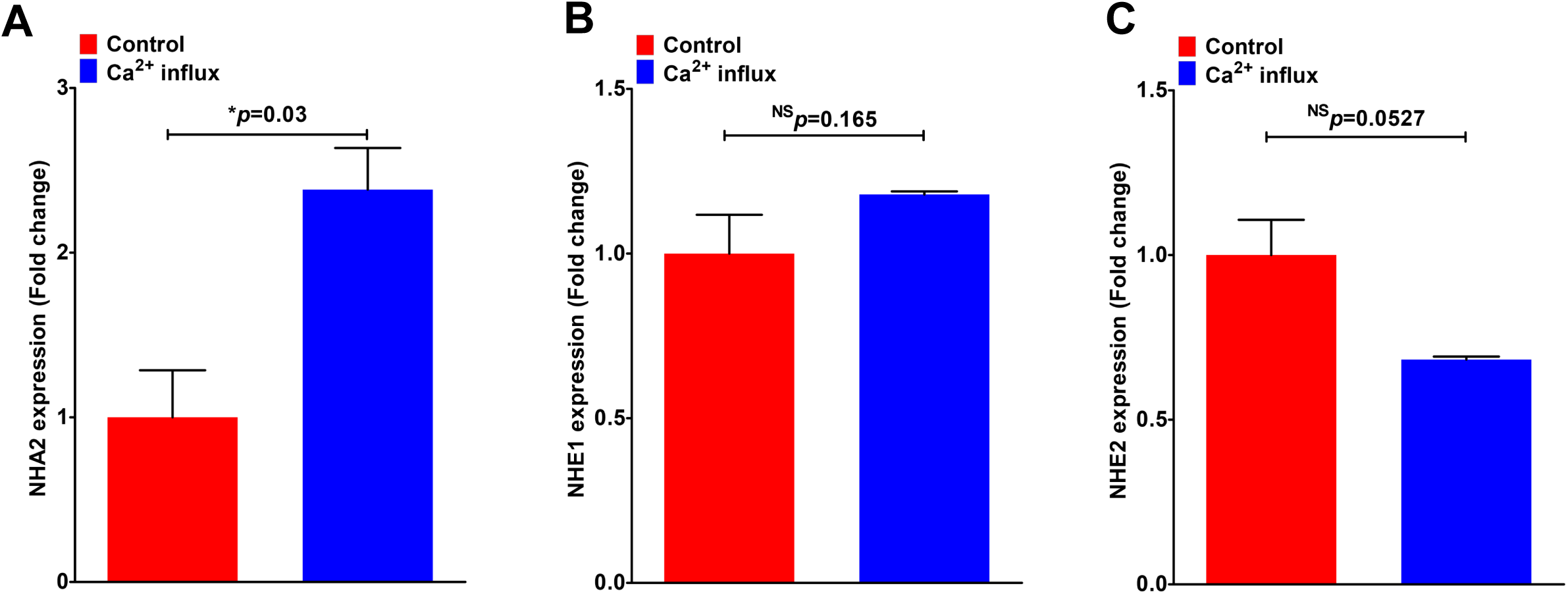
Ca^2+^-induced activation of NHA2 expression. A-C. Gene expression changes in response to Ca^2+^ influx following treatment with ionomycin and phorbol myristate in human primary T lymphoblasts obtained from the GSE50971 dataset. A. NHA2 showed significant up-regulation (~2.4-fold higher; Student’s t test; ^*^p=0.03). B. NHE1 expression showed no change (Student’s t test; p=0.165). C. NHE2 expression was lower, but non-significant (Student’s t test; p=0.0527).

**Fig. S3:**
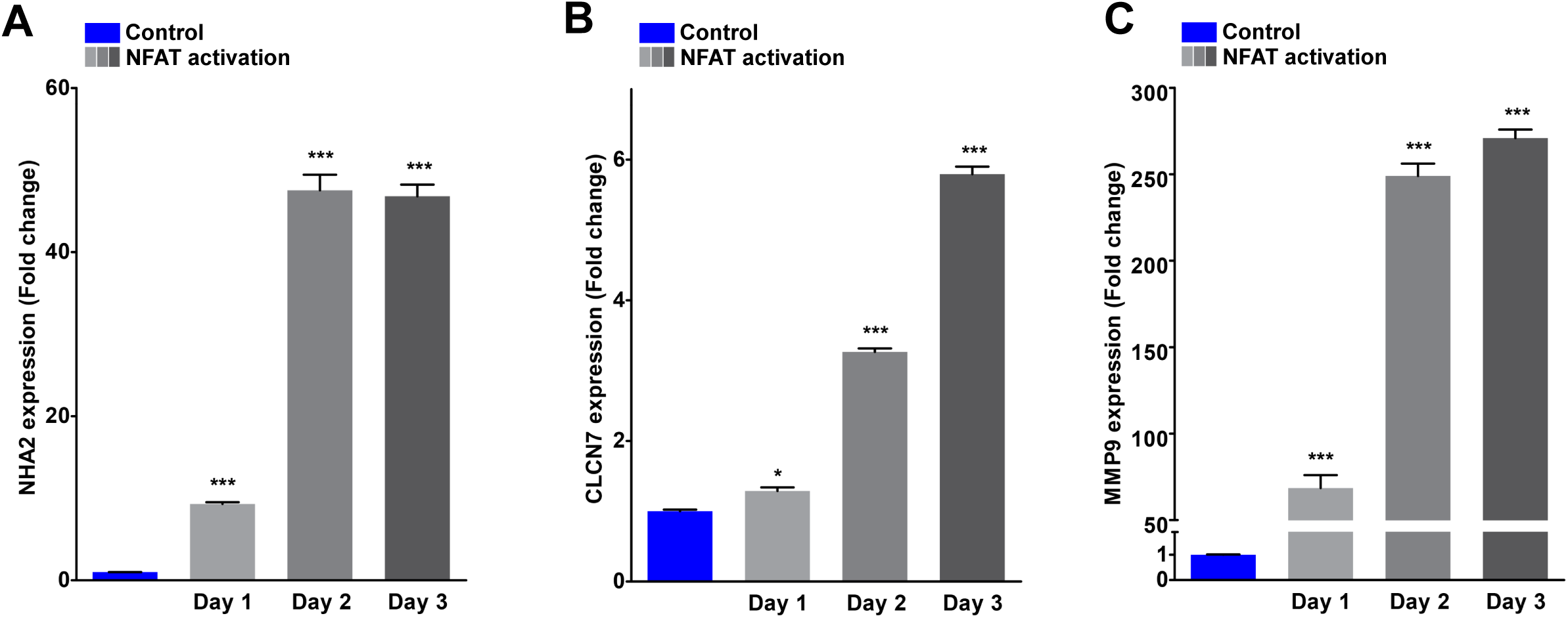
NFAT activation increases NHA2 expression. A-C. Gene expression changes were determined from the GSE57468 dataset. In this experiment, mouse bone marrow-derived osteoclast precursors were treated with RANKL for up to 3 days to differentiate into osteoclasts. A. Robust, time-dependent up-regulation of NHA2 levels in response to NFAT-mediated osteoclast differentiation *in vitro* (~10-fold on day 1; ~50-fold on day 2-3; Student’s t test; ^***^p<0.001). (B-C). Significant up-regulation of well-known NFAT target genes, CLCN7 (~1.3-fold on day 1; ~3.3-fold on day 2; ~5.8-fold on day 3; Student’s t test; ^*^p<0.05; ^***^p<0.001) and MMP9 (~69-fold on day 1; ~249-fold on day 2; ~270-fold on day 3; Student’s t test; ^***^p<0.001).

**Fig. S4:**
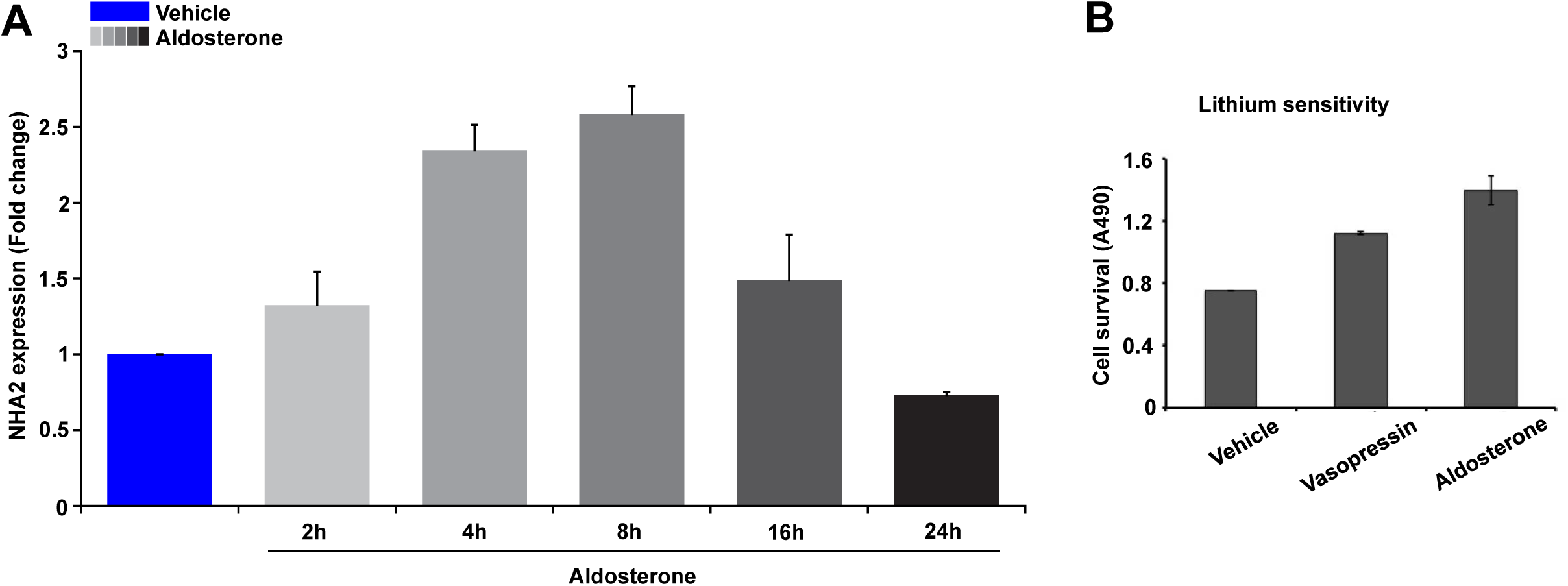
Hormonal induction of NHA2 expression. A. qPCR data showing fold change in NHA2 transcript levels following treatment of MDCK cells with aldosterone for indicated time periods ranging from 2-24h. Note significant and phasic increase in NHA2 expression with aldosterone treatment that peaked (~2.6-fold) at 8h and reached baseline by 24h. B. Lithium sensitivity assay to evaluate functional consequences of the hormonal induction of NHA2. Consistent with increased NHA2 mRNA and protein expression, treatment with either vasopressin or aldosterone for 8 h resulted in increased cell survival in media supplemented with 90 mM LiCl, relative to untreated control. Cell Survival in the presence of LiCl was measured using MTT assay.

**Fig. S5:**
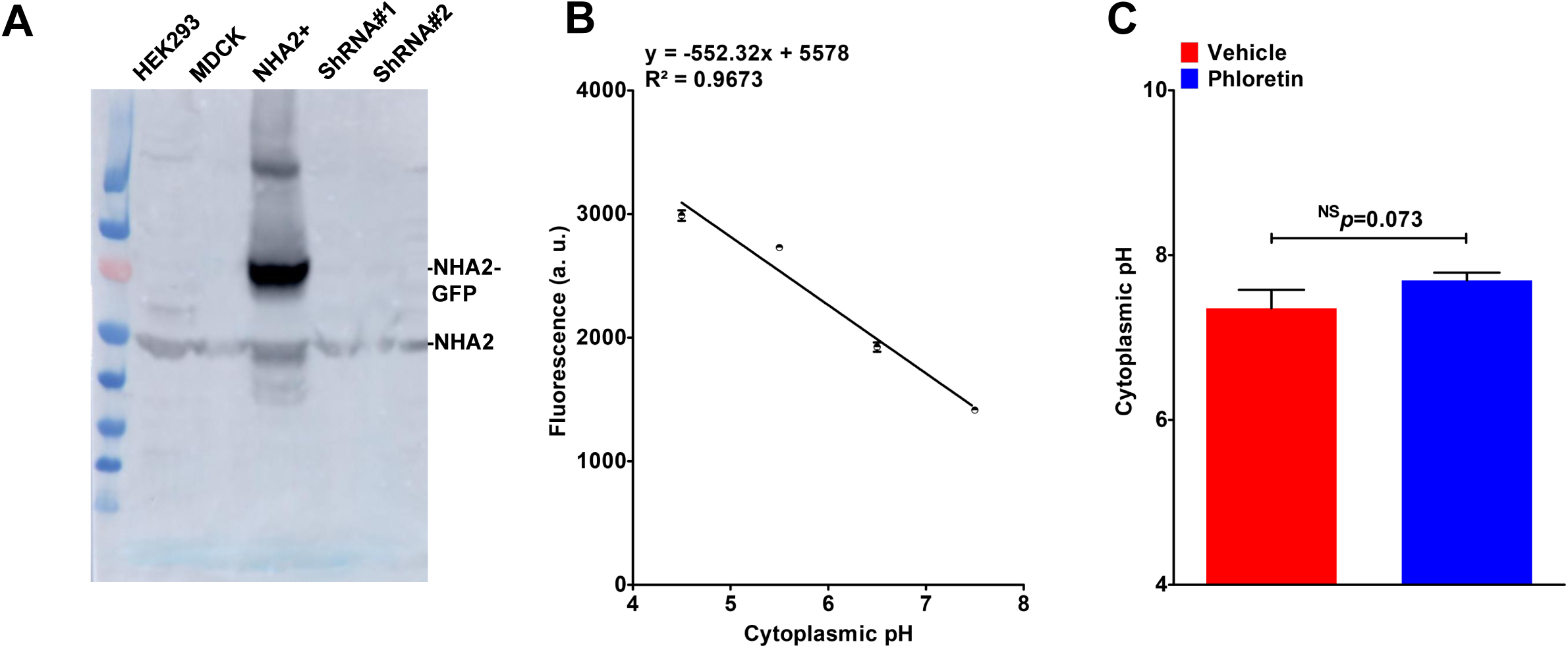
NHA2 inhibition does not alter cytoplasmic pH. A. Lentivirally mediated knockdown of NHA2 in NHA2+ MDCK cells, using two different lentiviral shRNA constructs (ShRNA#1 and ShRNA#2) targeting human NHA2, resulted in robust knockdown of GFP-tagged NHA2. Western blot of total protein was probed using anti-NHA2 antibody. (B-C). Cytoplasmic pH in NHA2+ cells with vehicle or phloretin treatment was determined, as described under Methods. Calibration curve is shown in B. Inhibition of NHA2 with phloretin did not significantly alter cytoplasmic pH (C) (n=3; Student’s t test; p=0.073).

## References

1. Ong, A. C., Devuyst, O., Knebelmann, B., Walz, G., and ERA-EDTA Working Group for Inherited Kidney Diseases. (2015) Autosomal dominant polycystic kidney disease: the changing face of clinical management. Lancet 385, 1993–2002

2. Harris, P. C., and Rossetti, S. (2010) Molecular diagnostics for autosomal dominant polycystic kidney disease. Nature reviews. Nephrology 6, 197–206

3. Hughes, J., Ward, C. J., Peral, B., Aspinwall, R., Clark, K., San Millan, J. L., Gamble, V., and Harris, P. C. (1995) The polycystic kidney disease 1 (PKD1) gene encodes a novel protein with multiple cell recognition domains. Nature genetics 10, 151–160

4. Nims, N., Vassmer, D., and Maser, R. L. (2003) Transmembrane domain analysis of polycystin-1, the product of the polycystic kidney disease-1 (PKD1) gene: evidence for 11 membrane-spanning domains. Biochemistry 42, 13035–13048

5. Qian, F., Germino, F. J., Cai, Y., Zhang, X., Somlo, S., and Germino, G. G. (1997) PKD1 interacts with PKD2 through a probable coiled-coil domain. Nature genetics 16, 179–183

6. Kim, H., Xu, H., Yao, Q., Li, W., Huang, Q., Outeda, P., Cebotaru, V., Chiaravalli, M., Boletta, A., Piontek, K., Germino, G. G., Weinman, E. J., Watnick, T., and Qian, F. (2014) Ciliary membrane proteins traffic through the Golgi via a Rabep1/GGA1/Arl3-dependent mechanism. Nature communications 5, 5482

7. Talbot, J. J., Shillingford, J. M., Vasanth, S., Doerr, N., Mukherjee, S., Kinter, M. T., Watnick, T., and Weimbs, T. (2011) Polycystin-1 regulates STAT activity by a dual mechanism. Proceedings of the National Academy of Sciences of the United States of America 108, 7985–7990

8. Olteanu, D., Liu, X., Liu, W., Roper, V. C., Sharma, N., Yoder, B. K., Satlin, L. M., Schwiebert, E. M., and Bevensee, M. O. (2012) Increased Na+/H+ exchanger activity on the apical surface of a cilium-deficient cortical collecting duct principal cell model of polycystic kidney disease. American journal of physiology. Cell physiology 302, C1436–1451

9. Avner, E. D., McDonough, A. A., and Sweeney, W. E., Jr. (2012) Transport, cilia, and PKD: must we in (cyst) on interrelationships? Focus on “Increased Na+/H+ exchanger activity on the apical surface of a cilium-deficient cortical collecting duct principal cell model of polycystic kidney disease”. American journal of physiology. Cell physiology 302, C1434–1435

10. Coaxum, S. D., Blanton, M. G., Joyner, A., Akter, T., Bell, P. D., Luttrell, L. M., Raymond, J. R., Sr., Lee, M. H., Blichmann, P. A., Garnovskaya, M. N., and Saigusa, T. (2014) Epidermal growth factor-induced proliferation of collecting duct cells from Oak Ridge polycystic kidney mice involves activation of Na+/H+ exchanger. American journal of physiology. Cell physiology 307, C554–560

11. Fuster, D. G., and Alexander, R. T. (2014) Traditional and emerging roles for the SLC9 Na+/H+ exchangers. Pflugers Archiv: European journal of physiology 466, 61–76

12. Donowitz, M., Ming Tse, C., and Fuster, D. (2013) SLC9/NHE gene family, a plasma membrane and organellar family of Na(+)/H(+) exchangers. Molecular aspects of medicine 34, 236–251

13. Kondapalli, K. C., Prasad, H., and Rao, R. (2014) An inside job: how endosomal Na(+)/H(+) exchangers link to autism and neurological disease. Frontiers in cellular neuroscience 8, 172

14. Xiang, M., Feng, M., Muend, S., and Rao, R. (2007) A human Na+/H+ antiporter sharing evolutionary origins with bacterial NhaA may be a candidate gene for essential hypertension. Proceedings of the National Academy of Sciences of the United States of America 104, 18677–18681

15. Fuster, D. G., Zhang, J., Shi, M., Bobulescu, I. A., Andersson, S., and Moe, O. W. (2008) Characterization of the sodium/hydrogen exchanger NHA2. Journal of the American Society of Nephrology: JASN 19, 1547–1556

16. Chen, S. R., Chen, M., Deng, S. L., Hao, X. X., Wang, X. X., and Liu, Y. X. (2016) Sodium-hydrogen exchanger NHA1 and NHA2 control sperm motility and male fertility. Cell death & disease 7, e2152

17. Kondapalli, K. C., Kallay, L. M., Muszelik, M., and Rao, R. (2012) Unconventional chemiosmotic coupling of NHA2, a mammalian Na+/H+ antiporter, to a plasma membrane H+ gradient. The Journal of biological chemistry 287, 36239–36250

18. Schushan, M., Xiang, M., Bogomiakov, P., Padan, E., Rao, R., and Ben-Tal, N. (2010) Model-guided mutagenesis drives functional studies of human NHA2, implicated in hypertension. Journal of molecular biology 396, 1181–1196

19. Kondapalli, K. C., Todd Alexander, R., Pluznick, J. L., and Rao, R. (2017) NHA2 is expressed in distal nephron and regulated by dietary sodium. Journal of physiology and biochemistry 73, 199–205

20. Song, X., Di Giovanni, V., He, N., Wang, K., Ingram, A., Rosenblum, N. D., and Pei, Y. (2009) Systems biology of autosomal dominant polycystic kidney disease (ADPKD): computational identification of gene expression pathways and integrated regulatory networks. Human molecular genetics 18, 2328–2343

21. Yang, B., Sonawane, N. D., Zhao, D., Somlo, S., and Verkman, A. S. (2008) Small-molecule CFTR inhibitors slow cyst growth in polycystic kidney disease. Journal of the American Society of Nephrology: JASN 19, 1300–1310

22. Sun, Y., Zhou, H., and Yang, B. X. (2011) Drug discovery for polycystic kidney disease. Acta pharmacologica Sinica 32, 805–816

23. Cebotaru, V., Cebotaru, L., Kim, H., Chiaravalli, M., Boletta, A., Qian, F., and Guggino, W. B. (2014) Polycystin-1 negatively regulates Polycystin-2 expression via the aggresome/autophagosome pathway. The Journal of biological chemistry 289, 6404–6414

24. Cross, B. M., Breitwieser, G. E., Reinhardt, T. A., and Rao, R. (2014) Cellular calcium dynamics in lactation and breast cancer: from physiology to pathology. American journal of physiology. Cell physiology 306, C515–526

25. Feng, M., Grice, D. M., Faddy, H. M., Nguyen, N., Leitch, S., Wang, Y., Muend, S., Kenny, P. A., Sukumar, S., Roberts-Thomson, S. J., Monteith, G. R., and Rao, R. (2010) Store-independent activation of Orai1 by SPCA2 in mammary tumors. Cell 143, 84–98

26. Woodward, O. M., Li, Y., Yu, S., Greenwell, P., Wodarczyk, C., Boletta, A., Guggino, W. B., and Qian, F. (2010) Identification of a polycystin-1 cleavage product, P100, that regulates store operated Ca entry through interactions with STIM1. PloS one 5, e12305

27. Villarroya-Beltri, C., Gutierrez-Vazquez, C., Sanchez-Cabo, F., Perez-Hernandez, D., Vazquez, J., Martin-Cofreces, N., Martinez-Herrera, D. J., Pascual-Montano, A., Mittelbrunn, M., and Sanchez-Madrid, F. (2013) Sumoylated hnRNPA2B1 controls the sorting of miRNAs into exosomes through binding to specific motifs. Nature communications 4, 2980

28. Pan, M. G., Xiong, Y., and Chen, F. (2013) NFAT gene family in inflammation and cancer. Current molecular medicine 13, 543–554

29. Kim, J. H., and Kim, N. (2014) Regulation of NFATc1 in Osteoclast Differentiation. Journal of bone metabolism 21, 233–241

30. Charles, J. F., Coury, F., Sulyanto, R., Sitara, D., Wu, J., Brady, N., Tsang, K., Sigrist, K., Tollefsen, D. M., He, L., Storm, D., and Aliprantis, A. O. (2012) The collection of NFATc1-dependent transcripts in the osteoclast includes numerous genes non-essential to physiologic bone resorption. Bone 51, 902–912

31. An, D., Kim, K., and Lu, W. (2014) Defective entry into mitosis 1 (Dim1) negatively regulates osteoclastogenesis by inhibiting the expression of nuclear factor of activated T-cells, cytoplasmic, calcineurin-dependent 1 (NFATc1). The Journal of biological chemistry 289, 24366–24373

32. Aliprantis, A. O., Ueki, Y., Sulyanto, R., Park, A., Sigrist, K. S., Sharma, S. M., Ostrowski, M. C., Olsen, B. R., and Glimcher, L. H. (2008) NFATc1 in mice represses osteoprotegerin during osteoclastogenesis and dissociates systemic osteopenia from inflammation in cherubism. The Journal of clinical investigation 118, 3775–3789

33. De Vries, T. J., Schoenmaker, T., Aerts, D., Grevers, L. C., Souza, P. P., Nazmi, K., van de Wiel, M., Ylstra, B., Lent, P. L., Leenen, P. J., and Everts, V. (2015) M-CSF priming of osteoclast precursors can cause osteoclastogenesis-insensitivity, which can be prevented and overcome on bone. Journal of cellular physiology 230, 210–225

34. Martinez, G. J., Pereira, R. M., Aijo, T., Kim, E. Y., Marangoni, F., Pipkin, M. E., Togher, S., Heissmeyer, V., Zhang, Y. C., Crotty, S., Lamperti, E. D., Ansel, K. M., Mempel, T. R., Lahdesmaki, H., Hogan, P. G., and Rao, A. (2015) The transcription factor NFAT promotes exhaustion of activated CD8(+) T cells. Immunity 42, 265–278

35. Blomberg, K. E., Boucheron, N., Lindvall, J. M., Yu, L., Raberger, J., Berglof, A., Ellmeier, W., and Smith, C. E. (2009) Transcriptional signatures of Itk-deficient CD3+, CD4+ and CD8+ T-cells. BMC genomics 10, 233

36. Knyazev, E. N., Mal’tseva, D. V., Zacharyants, A. A., Zakharova, G. S., Zhidkova, O. V., and Poloznikov, A. A. (2018) TNFalpha-Induced Expression of Transport Protein Genes in HUVEC Cells Is Associated with Enhanced Expression of Transcription Factor Genes RELB and NFKB2 of the Non-Canonical NF-kappaB Pathway. Bulletin of experimental biology and medicine 164, 757–761

37. Burn, S. F., Webb, A., Berry, R. L., Davies, J. A., Ferrer-Vaquer, A., Hadjantonakis, A. K., Hastie, N. D., and Hohenstein, P. (2011) Calcium/NFAT signalling promotes early nephrogenesis. Developmental biology 352, 288–298

38. Aguiari, G., Trimi, V., Bogo, M., Mangolini, A., Szabadkai, G., Pinton, P., Witzgall, R., Harris, P. C., Borea, P. A., Rizzuto, R., and del Senno, L. (2008) Novel role for polycystin-1 in modulating cell proliferation through calcium oscillations in kidney cells. Cell proliferation 41, 554–573

39. Puri, S., Magenheimer, B. S., Maser, R. L., Ryan, E. M., Zien, C. A., Walker, D. D., Wallace, D. P., Hempson, S. J., and Calvet, J. P. (2004) Polycystin-1 activates the calcineurin/NFAT (nuclear factor of activated T-cells) signaling pathway. The Journal of biological chemistry 279, 55455–55464

40. Arnould, T., Kim, E., Tsiokas, L., Jochimsen, F., Gruning, W., Chang, J. D., and Walz, G. (1998) The polycystic kidney disease 1 gene product mediates protein kinase C alpha-dependent and c-Jun N-terminal kinase-dependent activation of the transcription factor AP-1. The Journal of biological chemistry 273, 6013–6018

41. Kim, E., Arnould, T., Sellin, L. K., Benzing, T., Fan, M. J., Gruning, W., Sokol, S. Y., Drummond, I., and Walz, G. (1999) The polycystic kidney disease 1 gene product modulates Wnt signaling. The Journal of biological chemistry 274, 4947–4953

42. Prasad, H., and Rao, R. (2018) Histone deacetylase-mediated regulation of endolysosomal pH. The Journal of biological chemistry 293, 6721–6735

43. Belibi, F. A., Wallace, D. P., Yamaguchi, T., Christensen, M., Reif, G., and Grantham, J. J. (2002) The effect of caffeine on renal epithelial cells from patients with autosomal dominant polycystic kidney disease. Journal of the American Society of Nephrology: JASN 13, 2723–2729

44. Kong, H., Jones, P. P., Koop, A., Zhang, L., Duff, H. J., and Chen, S. R. (2008) Caffeine induces Ca2+ release by reducing the threshold for luminal Ca2+ activation of the ryanodine receptor. The Biochemical journal 414, 441–452

45. Nilsson, M., Hansson, E., and Ronnback, L. (1992) Agonist-evoked Ca2+ transients in primary astroglial cultures‐‐modulatory effects of valproic acid. Glia 5, 201–209

46. Suwanjang, W., Holmstrom, K. M., Chetsawang, B., and Abramov, A. Y. (2013) Glucocorticoids reduce intracellular calcium concentration and protects neurons against glutamate toxicity. Cell calcium 53, 256–263

47. Cao, Y., Semanchik, N., Lee, S. H., Somlo, S., Barbano, P. E., Coifman, R., and Sun, Z. (2009) Chemical modifier screen identifies HDAC inhibitors as suppressors of PKD models. Proceedings of the National Academy of Sciences of the United States of America 106, 21819–21824

48. Perry, P. J., Calloway, R. A., Cook, B. L., and Smith, R. E. (1984) Theophylline precipitated alterations of lithium clearance. Acta psychiatrica Scandinavica 69, 528–537

49. Grandjean, E. M., and Aubry, J. M. (2009) Lithium: updated human knowledge using an evidence-based approach: part III: clinical safety. CNS drugs 23, 397–418

50. Chebib, F. T., Sussman, C. R., Wang, X., Harris, P. C., and Torres, V. E. (2015) Vasopressin and disruption of calcium signalling in polycystic kidney disease. Nature reviews. Nephrology 11, 451–464

51. Stewart, G. S., Thistlethwaite, A., Lees, H., Cooper, G. J., and Smith, C. (2009) Vasopressin regulation of the renal UT-A3 urea transporter. American journal of physiology. Renal physiology 296, F642–648

52. Carmosino, M., Rizzo, F., Ferrari, P., Torielli, L., Ferrandi, M., Bianchi, G., Svelto, M., and Valenti, G. (2011) NKCC2 is activated in Milan hypertensive rats contributing to the maintenance of salt-sensitive hypertension. Pflugers Archiv : European journal of physiology 462, 281–291

53. Stiber, J. A., Tabatabaei, N., Hawkins, A. F., Hawke, T., Worley, P. F., Williams, R. S., and Rosenberg, P. (2005) Homer modulates NFAT-dependent signaling during muscle differentiation. Developmental biology 287, 213–224

54. Scicchitano, B. M., Spath, L., Musaro, A., Molinaro, M., Rosenthal, N., Nervi, C., and Adamo, S. (2005) Vasopressin-dependent myogenic cell differentiation is mediated by both Ca2+/calmodulin-dependent kinase and calcineurin pathways. Molecular biology of the cell 16, 3632–3641

55. Kjeldsen, S. E., Os, I., Forsberg, G., Aakesson, I., Skjoto, J., Frederichsen, P., Fonstelien, E., and Eide, I. (1985) Dietary sodium intake increases vasopressin secretion in man. Journal of clinical hypertension 1, 123–131

56. Levine, S., Franki, N., and Hays, R. M. (1973) Effect of phloretin on water and solute movement in the toad bladder. The Journal of clinical investigation 52, 1435–1442

57. Chapman, A. B., Johnson, A., Gabow, P. A., and Schrier, R. W. (1990) The renin-angiotensin-aldosterone system and autosomal dominant polycystic kidney disease. The New England journal of medicine 323, 1091–1096

58. Tkachenko, O., Helal, I., Shchekochikhin, D., and Schrier, R. W. (2013) Renin-Angiotensin-aldosterone system in autosomal dominant polycystic kidney disease. Current hypertension reviews 9, 12–20

59. Hasler, U., Mordasini, D., Bianchi, M., Vandewalle, A., Feraille, E., and Martin, P. Y. (2003) Dual influence of aldosterone on AQP2 expression in cultured renal collecting duct principal cells. The Journal of biological chemistry 278, 21639–21648

60. Perlewitz, A., Nafz, B., Skalweit, A., Fahling, M., Persson, P. B., and Thiele, B. J. (2010) Aldosterone and vasopressin affect {alpha}- and {gamma}-ENaC mRNA translation. Nucleic acids research 38, 5746–5760

61. Coutry, N., Farman, N., Bonvalet, J. P., and Blot-Chabaud, M. (1995) Synergistic action of vasopressin and aldosterone on basolateral Na(+)-K(+)-ATPase in the cortical collecting duct. The Journal of membrane biology 145, 99–106

62. Schafer, J. A., and Hawk, C. T. (1992) Regulation of Na+ channels in the cortical collecting duct by AVP and mineralocorticoids. Kidney international 41, 255–268

63. de Almeida, P. W., de Freitas Lima, R., de Morais Gomes, E. R., Rocha-Resende, C., Roman-Campos, D., Gondim, A. N., Gavioli, M., Lara, A., Parreira, A., de Azevedo Nunes, S. L., Alves, M. N., Santos, S. L., Alenina, N., Bader, M., Resende, R. R., dos Santos Cruz, J., Souza dos Santos, R. A., and Guatimosim, S. (2013) Functional cross-talk between aldosterone and angiotensin-(1-7) in ventricular myocytes. Hypertension 61, 425–430

64. Misfeldt, D. S., Hamamoto, S. T., and Pitelka, D. R. (1976) Transepithelial transport in cell culture. Proceedings of the National Academy of Sciences of the United States of America 73, 1212–1216

65. Lever, J. E. (1979) Inducers of mammalian cell differentiation stimulate dome formation in a differentiated kidney epithelial cell line (MDCK). Proceedings of the National Academy of Sciences of the United States of America 76, 1323–1327

66. Su, H. W., Yeh, H. H., Wang, S. W., Shen, M. R., Chen, T. L., Kiela, P. R., Ghishan, F. K., and Tang, M. J. (2007) Cell confluence-induced activation of signal transducer and activator of transcription-3 (Stat3) triggers epithelial dome formation via augmentation of sodium hydrogen exchanger-3 (NHE3) expression. The Journal of biological chemistry 282, 9883–9894

67. N, O. B., Husson, H., Dackowski, W. R., Lawrence, B. D., Clow, P. A., Roberts, B. L., Klinger, K. W., and Ibraghimov-Beskrovnaya, O. (2002) Functional polycystin-1 expression is developmentally regulated during epithelial morphogenesis in vitro: down-regulation and loss of membrane localization during cystogenesis. Human molecular genetics 11, 923–936

68. Rodrig, N., Osanai, T., Iwamori, M., and Nagai, Y. (1988) Uncoupling of intracellular cyclic AMP and dome formation in cultured canine kidney epithelial cells: effects of gangliosides and vasopressin. Journal of biochemistry 104, 215–219

69. Oberleithner, H., Vogel, U., and Kersting, U. (1990) Madin-Darby canine kidney cells. I. Aldosterone-induced domes and their evaluation as a model system. Pflugers Archiv: European journal of physiology 416, 526–532

70. Terryn, S., Ho, A., Beauwens, R., and Devuyst, O. (2011) Fluid transport and cystogenesis in autosomal dominant polycystic kidney disease. Biochimica et biophysica acta 1812, 1314–1321

71. Grantham, J. J., Chapman, A. B., and Torres, V. E. (2006) Volume progression in autosomal dominant polycystic kidney disease: the major factor determining clinical outcomes. Clinical journal of the American Society of Nephrology : CJASN 1, 148–157

72. Vareesangthip, K., Thomas, T. H., and Wilkinson, R. (1995) Abnormal effect of thiol groups on erythrocyte Na/Li countertransport kinetics in adult polycystic kidney disease. Nephrology, dialysis, transplantation : official publication of the European Dialysis and Transplant Association - European Renal Association 10, 2219–2223

73. Shah, S., Watnick, T., and Atta, M. G. (2010) Not all renal cysts are created equal. Lancet 376, 1024

74. Kling, M. A., Fox, J. G., Johnston, S. M., Tolkoff-Rubin, N. E., Rubin, R. H., and Colvin, R. B. (1984) Effects of long-term lithium administration on renal structure and function in rats. A distinctive tubular lesion. Laboratory investigation; a journal of technical methods and pathology 50, 526–535

75. Liu, H. M., He, J. Y., Zhang, Q., Lv, W. Q., Xia, X., Sun, C. Q., Zhang, W. D., and Deng, H. W. (2018) Improved detection of genetic loci in estimated glomerular filtration rate and type 2 diabetes using a pleiotropic cFDR method. Molecular genetics and genomics : MGG 293, 225–235

